# A quantitative approach to easily characterize and predict cell death and escape to β-lactam treatments

**DOI:** 10.1101/2021.07.16.452741

**Authors:** Virgile Andreani, Viktoriia Gross, Lingchong You, Philippe Glaser, Imane El Meouche, Gregory Batt

## Abstract

Commensal and pathogenic *E. coli* strains are increasingly found to be resistant to β-lactams, one of the most widely prescribed classes of antibiotics. Understanding escape to such treatments is complex since β-lactams have several cellular targets and since several mechanisms might be involved in treatment escape in a combined manner. Surprisingly, the accumulated knowledge has not yet proven effective enough to predict the bacterial response to antibiotic treatments at both cellular and population levels with quantitative accuracy for -producing bacteria. Here, we propose a mathematical model that captures in a comprehensive way key phenomena happening at the molecular, cellular, and population levels, as well as their interactions. Our growth-fragmentation model gives a central role to cellular heterogeneity and filamentation as a way for cells to gain time until the degradation of the antibiotic by the β-lactamases released by the dead cell population. Importantly, the model can account for the observed temporal evolution of the total (live and dead) biomass and of the live cell numbers for various antibiotic concentrations. To our knowledge, this is the first model able to quantitatively reconciliate these two classical views on cell death (OD and CFUs) for clinical isolates expressing extended-spectrum beta-lactamases (ESBL). Moreover, our model has a strong predictive power. When calibrated using a slight extension of OD-based data that we propose here, it can predict the CFU profiles in initial and delayed treatments despite inoculum effects, and suggest non-trivial optimal treatments. Generating quality data in quantity has been essential for model development and validation on non-model E. coli strains. We developed protocols to increase the reproducibility of growth kinetics assays and to increase the throughput of CFU assays.

## Introduction

*Escherichia coli* is a major cause of many common infections, including urinary tract infections affecting over 150 million people worldwide, primarily women^1^. An increasing number of commensal and pathogenic *E. coli* strains are found to be resistant to β-lactams, one of the most widely prescribed classes of antibiotics^2,3^.

Understanding escape to β-lactam treatments is complex since β-lactams might have several cellular targets and since several mechanisms might be involved in treatment escape in a combined manner^4,5^. Regarding targets, these penicillin-derived antibiotics disable the capacity of cells to synthesize and maintain their cell wall and prevent proper cell septation, in both cases by inactivating essential enzymes involved in peptidoglycan biosynthesis, known as penicillin-binding proteins (PBP1 and PBP3 notably). Regarding escape mechanisms, the most important one is the expression of β-lactamases, enzymes that degrade the antibiotic. These enzymes are active both inside (periplasm) and outside (environment) the cell. Consequently, they provide a direct protection to the cell and an indirect protection to the cell as a member of a population. Indeed, the death of a part of the cell population releases β-lactamases in the environment, which degrade the antibiotic and facilitate the survival of the rest of the population. This phenomena, known as collective antibiotic tolerance (CAT) is at the origin of the well-known inoculum effect by which dense cell populations appear more resistant than dilute cell cultures^6^. Other important and commonly encountered escape mechanisms are the mutation of PBPs, the targets of the antibiotic, and the alteration of β-lactams influx and efflux in the cell via mutations of porin proteins or modification of efflux pumps. Overall, the capacity of bacterial cells to escape β-lactam treatments results from a complex combination of multiple factors.

A model-based approach is needed to capture these processes in a quantitative manner. However, the calibration of such a model would be a challenge. The gold standard approach to measure the temporal evolution of the live cell number upon antibiotic treatments is based on colony-forming unit (CFU) assays. These assays are used to count the number of live cells in a sample and consist in performing serial dilutions of the culture, in spreading small volumes of these dilutions on agar plates, and, after incubation, in counting the number of resulting colonies, indicating the initial number of live cells in the sample. CFU assays are highly informative and extremely simple in principle, but very labor intensive when the numbers of conditions and time points are large. A possible alternative is the use of growth kinetics assays. They consist in measuring at regular time intervals the optical density (OD) of a bacterial culture subjected to an antibiotic treatment. Many conditions can be followed in parallel thanks to the use of 96-well plates and plate-readers. This method is comparatively much easier. However, it suffers from two major limitations regarding the measurement of live cell numbers. Firstly, the measured OD is representative of the total biomass, both live and dead. Secondly, biomass and cell numbers are decorrelated in presence of significant cell filamentation. Because of their mechanisms of action, the latter is often encountered in β-lactam treatments. Because of the complexity of the processes and of the lack of quality quantitative data, no consolidated vision and no model have been proposed so far that can quantitatively explain the escape of a bacterial cell population to these widely-used treatments.

Here, we propose a model that captures in a comprehensive way key phenomena happening at the molecular, cell physiology, and population levels, as well as their interactions. Notably, the model gives a central role to filamentation as a way for cells to gain time until the degradation of the antibiotic by the released β-lactamases. Importantly, the model can account for the observed temporal evolution of the OD and live cell number for various clinical isolates and antibiotic concentrations of cefotaxime, as well as ceftriaxone and mecillinam. To our knowledge, this is the first model able to reconciliate these two classical views on cell death for clinical isolates expressing β-lactamases. We also provide an approach for its effective calibration based on a simple extension of growth kinetics assays, using repeated treatments. The model is then able to predict the temporal evolution of live cell numbers (CFU assays) solely based on OD readouts. This has significant practical interest given the laborious aspect of CFU assays. Lastly, our model is able to predict optimal treatments in non-trivial cases. To conduct this study, the ability to generate quality data in a rather large quantity for non-model *E. coli* strains was key. In this work, we also propose protocols to increase the reproducibility of growth kinetics assays and to increase the throughput of CFU assays.

## Results

### Experimental setup and modeling assumptions

For this study we selected 11 clinical isolates with different origins, different genetic backgrounds, and different levels of antibiotic susceptibility^7,8^. These strains express a panel of different families of β-lactamases, including carbapenemases, some strains expressing several of them simultaneously. Some of these strains also contain mutations in genes coding for proteins known for influencing antibiotic response, such as porins (*ompC, ompF*), PBP3 (ftsI), or DNA gyrase and topoisomerase subunits (*gyrA* and *parC* respectively). This selection contains strains from both human and animal infections, represents several sequence types and ranges from highly susceptible to highly resistant to β-lactam treatments. Full description of the collection is in Supplementary Text 1. We mainly study cefotaxime treatments. Cefotaxime is a broad spectrum third generation cephalosporin. We also provide results for ceftriaxone and mecillinam treatments.

At the center of our work, we focus on two simple questions. Can we reconciliate the growth kinetics assay view and the CFU assay view on cell death (Q1)? Can we predict CFUs from growth kinetics measurements (Q2). These questions appear to be very simple, but question at a fundamental level our understanding of β-lactam treatment escape (Fig 1a). To investigate these questions in a quantitative manner, we introduce a mechanistic model capturing key processes. On Fig1b, we propose a typical observation, an interpretation of the main processes underlying treatment escape, consistent with generally accepted knowledge, and a translation in terms of modeling assumptions. During the first hours following the addition of a significant dose of β-lactams, an exponential increase in OD can be observed, similarly to the control without antibiotics, but the number of live cells does not (it can even decrease). Cells do not divide but keep growing in biomass: they filament. Following this first phase, the OD of the treated population stays steady or slightly decreases for a few hours, while the number of live cells, indicated by the CFU curve, crashes: overly long filamented cells do not sustain constant biomass increase. After some time, the number of live cells transitions from rapid decrease to rapid increase: the antibiotic has been cleared from the culture medium by the β-lactamases released by lysed cells. After some additional time, regrowth starts to be detectable through OD measurements, and we can observe an increase in both OD and CFUs. Note that the moment of antibiotic clearance, a key event to understand the overall dynamics, is not visible in growth kinetics assays. In case of treatments with higher antibiotic concentrations (light pink line), the degradation of the antibiotic might be not sufficiently fast for the population to restart growth in the observation time frame.

**Figure 1.**
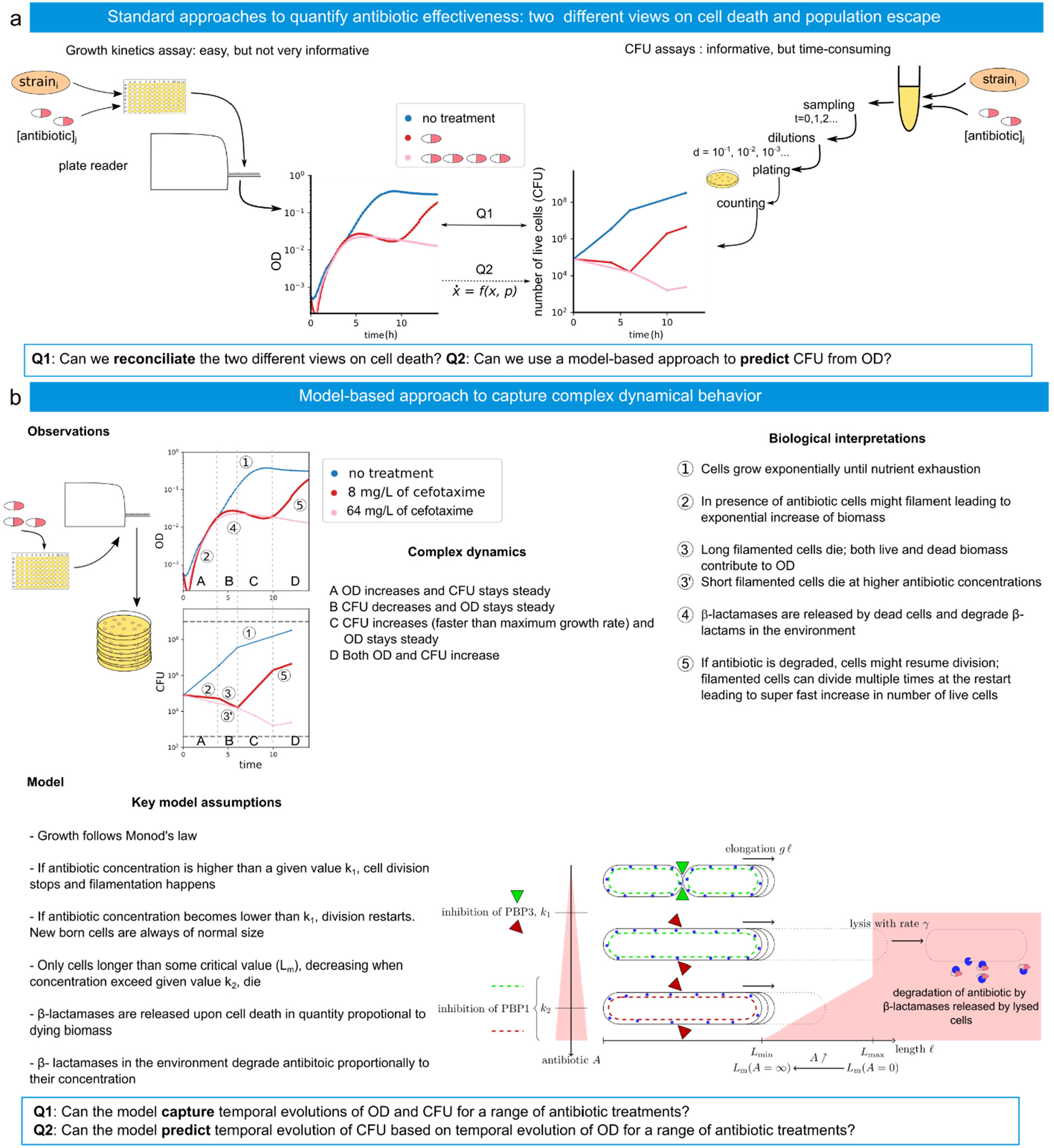
Characterizing and modeling the response of a bacterial population to β-lactam treatments. **a** Standard approaches to quantify antibiotic effectiveness correspond to growth kinetics assays, quantifying the temporal evolution of the biomass, and CFU assays, quantifying the temporal evolution of the live cell number. Growth kinetics assay*s* consists in following the optical density (OD) of liquid cultures inoculated with an isolate to be tested and treated with a given antibiotic concentration. CFU assays consists in taking samples of a liquid culture treated with antibiotics, in making serial dilutions of these samples, plating these dilutions on solid media, and after incubation, in counting the colonies of the media. The resulting dynamics are markedly different. **b** A model-based approach to capture the complex dynamics of cell death and population escape. The observation of a clinical isolate (here NILS56) treated with different concentrations of antibiotics shows a complex dynamics. These observations can be interpreted as following: 1) in the absence of antibiotics (blue line) cells grow exponentially until nutrient exhaustion; 2) in the presence of antibiotic (red line, A) cells might filament leading to exponential increase of biomass (OD) without increase in number (CFU); 3) long filamented cells die (B); both live and dead biomass contribute to OD; 3′) short filamented cells die at higher antibiotic concentrations (pink line); 4) β-lactamases are released by the lysed cells and degrade β-lactams in the environment; 5) if the antibiotic has been cleared, cells might resume division (C&D); filamented cells can divide multiple times at the restart leading to super-fast increase in live cell number. These interpretations leads to a set of modeling assumptions where the inhibition of PBP3 triggers filamentation, and the inhibition of PBP1 weakens the cell wall and facilitates the lysis of filamented cells, and the subsequent release of β-lactamases in the media. The population might regrow if the media is cleared from antibiotics before the death of all cells.

As evidenced here, there is a need for a model that captures various processes happening at different scales. At the cellular scale, one should notably take into account the fact that β-lactams target cell septation and cell wall maintenance in different ways (mainly targeting PBP3 and/or PBP1, and possibly at different concentration thresholds. It is known that for a given β-lactam concentration, the time to cell lysis is inversely proportional to the growth rate^9,10^. Because the elongation rate is not significantly affected by the antibiotic, this suggests that the death mechanism can be modelled with a critical cell length, reachable when cell septation is blocked^11^. Cell filaments longer than this threshold experience a significant death rate. This makes cell lysis a direct function not of the antibiotic itself but of cell length, that drastically increases because of antibiotic. The antibiotic can however, at high doses, reduce this critical length, through the inhibition of other PBPs such as the PBP1s which have a fundamental role in the repair of wall defects^12^. Moreover, simply considering that all the cells have the same length, which increases with time because of filamentation and eventually causes cell death, cannot explain the lysis of a subset only of the cell population. Only the longest cells die and this creates a non-trivial feedback on the average length of the surviving cells. Hence, accounting for the heterogeneity in cell lengths in the surviving cell population is required. Therefore, we developed a partial differential equation (PDE) model of the temporal evolution of the distribution of cell lengths within the population, and described with ordinary differential equations (ODEs) all other processes. The evolution of the cell length distribution is represented by a process of growth-fragmentation that relies on the assumption that cells elongate exponentially and divide with a rate depending on their length, into more than two fragments, which is notably the case for filamented cells after removal of the antibiotic^13^. The division rate is assumed to be a decreasing function of the antibiotic concentration in the medium, consistent with the observation that cefotaxime inhibits PBP3, involved in the septum formation^14,15^. Lastly, the cell elongation rate is assumed to be only a function of the nutrient concentration, and remains unchanged by the antibiotic, as is the case for most β-lactams.

At the macroscopic level, the evolution of the concentrations of nutrients, antibiotics and β-lactamases in the culture medium are described simple mechanistic ODEs. Some of our strains express several β-lactamases with different efficiencies. Yet, the model aggregates them into a single representative one. Lastly, the model also accounts for the fragments of lysed cells (the dead biomass) that may significantly contribute to the measured OD. The equations of the corresponding PDE/ODE model are given in the methods section and are described in more detail in the Supplementary Text 2.

To make the model computationally tractable, we eliminated the explicit representation of the cell length distribution *n*(*l, t*) to keep only its first moments: number of cells *N* and average length *L*. The ODEs for these quantities involve partial moments of the cell length distribution, which we managed to express only in terms of *N* and *L* through careful approximations (Supplementary Text 2). This leads us to our first question: can the model capture temporal evolution of OD and CFU for a range of antibiotic treatments?

### Q1: Can the model capture temporal evolutions of OD and CFU for a range of antibiotic treatments?

Our approach to investigate this question consists in exposing our 11 clinical isolates to 8 different concentrations of cefotaxime, measuring the temporal evolution of OD using plate-reader and the temporal evolution of CFU using droplet-based method, and calibrating the model on OD and CFU data (Figure 2a). More specifically, cells have been grown in 96-well microtiter plates in M9 minimal medium with 1 g/L glucose and 0.1% Tween 20, and the chosen antibiotic concentration has been applied at the beginning of the experiment. The OD has been followed using a Tecan Spark plate reader and samples have been regularly taken to perform CFU assays. As detailed in Supplementary Text 3, we observed for some isolates and in some conditions the presence of bacterial biofilms at the bottom of the wells. This was revealed through systematic measurements of the OD at different positions in microplate wells and through Crystal Violet staining at the end of experiments. This problem was a source of significant day-to-day variability in OD data. It was solved by the addition of a small quantity of Tween 20. Performing CFU assays for 96 conditions in parallel and with a 2-hour time resolution is highly impractical using standard approaches. This would require plating and counting more than 3000 Petri dishes per experiment. We therefore developed a protocol to perform CFU counts in a droplet-based manner that is conveniently performed using 96-tips pipettes and a handful of agar plates. We also adapted and retrained a software tool, CFU Spot Reader, to identify spots on plates, count colony numbers in spots and generate final counts and statistics for a given experiment. This is documented in Supplementary Text 4.

**Figure 2.**
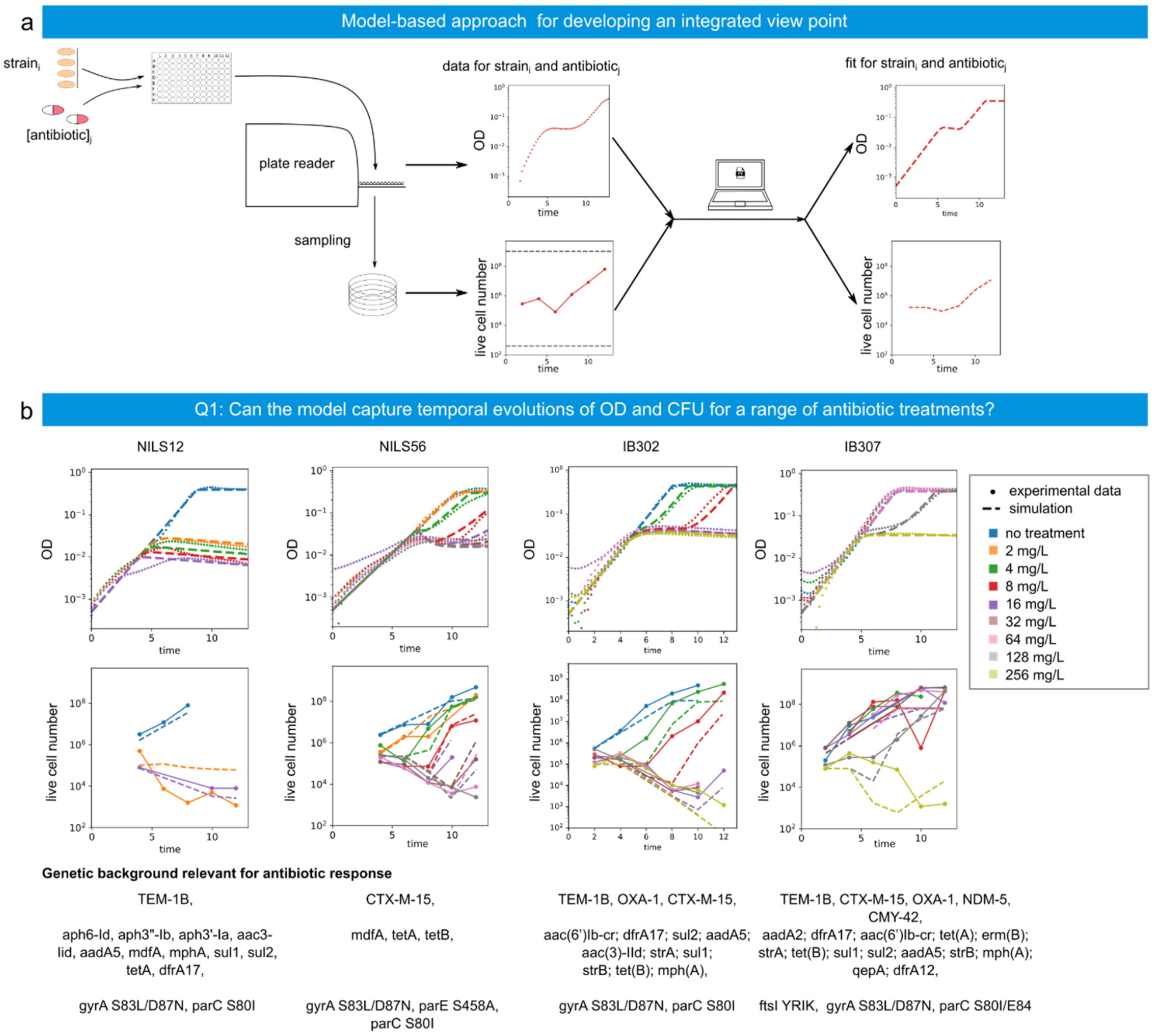
Characterizing and modeling the response of various clinical isolates to cefotaxime treatments. **a** A 96-well plate is inoculated with the isolates to be tested, and a range of antibiotic concentrations is applied. The OD is measured. Every two hours a sample is taken for CFU assays. The data from both OD and CFUs is then fed into a model calibration pipeline. Numerical parameters are estimated for each isolate and simulations corresponding to OD and CFU data are generated and compared to the experimental results. **b** Comparison of experimental data and the model fits. Four clinical isolates, representing a range of susceptibility to β-lactam treatment from very susceptible (NILS12), to highly resistant (IB307), are treated with different concentrations of cefotaxime, and optical density and live cell number are measured. Experimental data (points with (CFU) or without (OD) solid lines) and outputs of the model calibrated on OD and CFU data (dashed lines) show good agreements.

On figure 2b we present the results of characterization experiments for four strains. This data set has been used to calibrate the model, and model simulations are represented on Fig 2b as well. Data for all strains and the corresponding model simulations are presented in Supplementary Text 5. From OD data for the most susceptible strain, NILS12, we can observe that growth of biomass stops earlier at higher antibiotic concentrations, which we do not observe for other strains. From intermediate strains (for example, IB302, third column) we can observe that live cell number stays steady during the first hours while biomass increases exponentially which suggests presence of filamentation. Another interesting observation from the same data is that the death rate does not depend on antibiotic concentration, but the duration of cell death and hence the dead biomass does. These effects are well captured by our model and they go along with the main assumptions of the model such as filamentation, lysis caused by reaching a critical length and finally dependence of the critical length on antibiotic concentration. In the case of low concentration of antibiotic, given the low precision of CFU assays, the number of dead cells is not noticeable and dead cells are hidden by the start of the regrowth. This is the case of NILS56 in the presence of 4 mg/L, IB302 in the presence of 4 or 8 mg/L and IB307 in the presence of 128 mg/L. These cases being prone to high variability, they are difficult to capture and our model sometimes tends to overestimate the degree of cell lysis. Other interesting and not trivial observations come from looking into the first hours of the regrowth. Firstly, regrowth is initially hidden by dead biomass and, therefore, is not detectable from OD data, which can be observed from IB302 at 16 mg/L, NILS56 at 16, 32 or 64 mg/L. Secondly, in some cases (for example NILS56 at 8 mg/L) the number of live cells increase faster than the maximal growth rate, which can be explained by a de-filamentation effect, i.e. filaments dividing into more than 2 smaller cells. Despite their complexity and their limited observability, both of these phenomena are well captured by the model, which is impressive given their complexity and their limited observability.

Overall, our model offers a good agreement with the data for all 4 strains, therefore, proving its capacity to quantitatively capture temporal evolutions of both OD and CFUs despite the decorrelation between these two measurement modes, and to do so for a range of antibiotic treatments for a variety of clinical isolates with various cefotaxime sensitivity levels. Other kinetic models in the field describe only OD or only CFUs, and often in non-mechanistic ways, for example with an artificial delay after the start of the experiment to provoke the growth arrest. To our knowledge, our model is the first to reconcile these quantities, and to do so with mechanistic arguments.

### Q2: Can the model predict temporal evolution of CFU based on temporal evolution of OD for a range of antibiotic treatments?

To further challenge the predictive power of our model, we tested its capacity to predict the temporal evolution of live cell numbers based solely on OD data (Figure 3a). This was also motivated by the fact that, even using a droplet-based protocol, performing CFU assays remains a time-consuming and complicated task, especially when compared to the generation of OD curves by an automated plate-reader.

**Figure 3.**
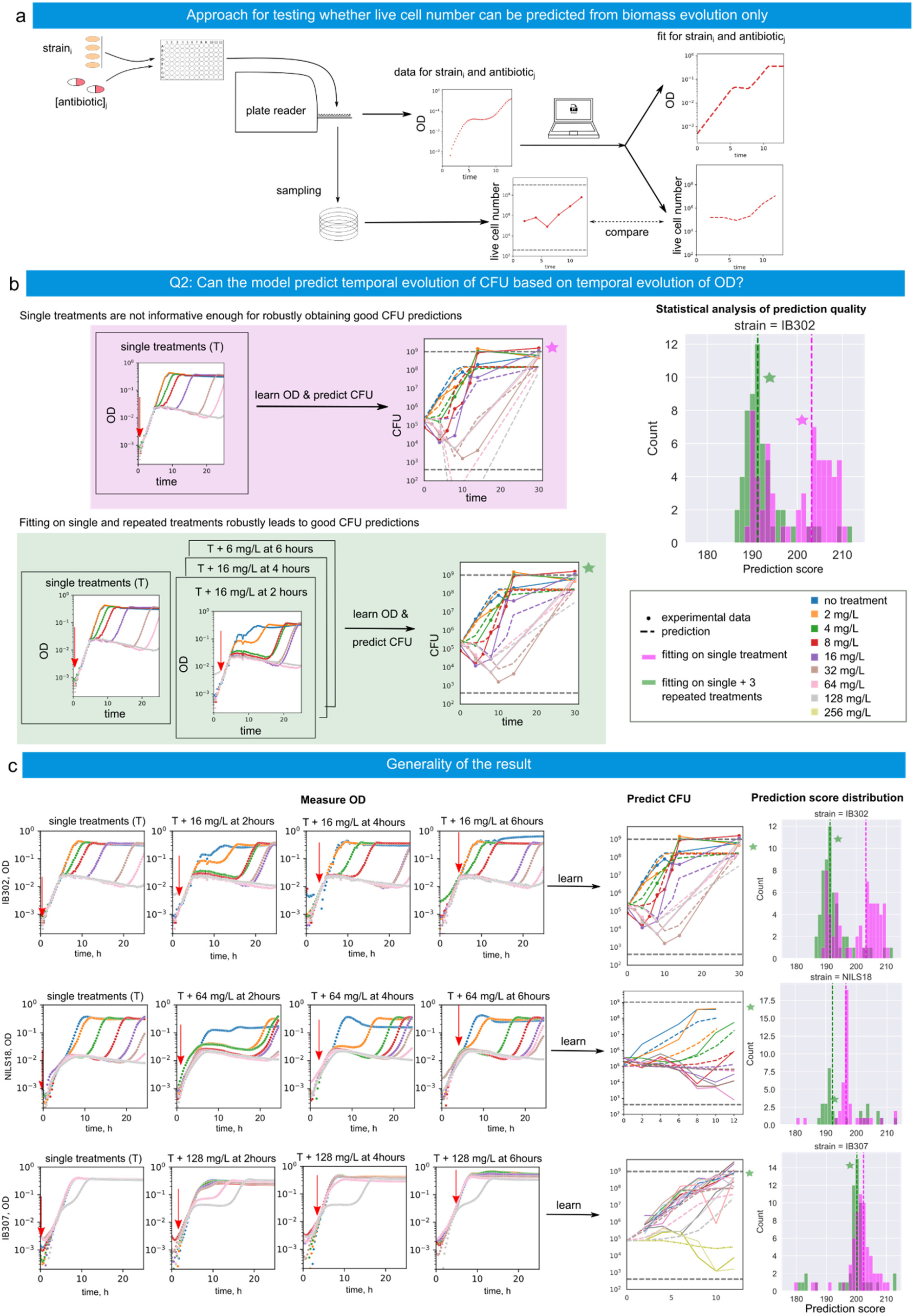
Prediction of the temporal evolution of CFUs based on the temporal evolution of OD. **a** 96-well plates are inoculated with the isolates to be tested, and a range of antibiotic concentrations is applied at the beginning (single treatments). For some experiments, a second antibiotic dose is applied several hours later (repeated treatments). The OD is read periodically, and samples are taken every two hours for CFU assays. The model is calibrated using OD data only. Simulations corresponding to both OD and CFU data are then performed and compared to the experimental results. **b** Parameter value estimates are obtained by repeatedly fitting a calibration dataset containing only an experiment with single treatments (top). For each parameter estimate, a CFU prediction score is calculated (histogram, magenta color). The CFU predictions shown on the left are generated with the parameter set having the median CFU predictions score. Other parameter value estimates are obtained by repeatedly fitting a dataset containing both single treatment experiments and three different repeated treatment experiments (bottom). The distribution of the corresponding CFU prediction scores is presented on the histogram (green color). The CFU predictions shown on the left are generated using the parameter set with the median prediction score. Experimental data is shown in points and solid lines, whereas model simulations are shown in dashed lines. **c** Generality of the result. Each row corresponds to one clinical isolate. For each isolate, we consider calibration datasets containing the OD measurements present either in only a single treatment experiment or in a single treatment experiment and in 3 repeated treatment experiments (left). The distribution of CFU prediction scores of different parameter value sets obtained by repeatedly fitting the model either using single treatments only (magenta) or using all the calibration data considered (green; right). The plot in the middle shows a representative prediction of the temporal evolution of CFUs using the set of parameter values having the median prediction score (center).

For one of the non-susceptible strains (IB302), we recalibrated the model, this time using only OD data, and we compared the predicted live cell numbers to the observed ones. For assessing the quality of the prediction, we repeated the fitting process forty times. Fitting is based on stochastic optimization tools and therefore generally returns different parameter sets, that generally have comparable fitting scores. For each of the forty different sets of parameter values, we calculated the CFU predictions score, defined as the mean square error of the logarithm of cell counts for all time points and all concentrations of antibiotic. The distribution of these scores is shown on Fig 3b (violet histogram), together with a typical CFU prediction, that is, a CFU prediction having a median quality. In all tested strains, it is possible within 40 fitting attempts to reach a parameter value set that accurately predicts the live cell number. However, this fitting process does not robustly lead to good predictions. Moreover, the parameter sets having best fitting scores (on OD data) are not those with best prediction score (on CFU data), indicating the presence of overfitting (Supplementary Text 8). This suggests that the data is not informative enough to sufficiently constrain model parameters to obtain a predictive model.

Therefore, we sought to increase the informativeness of our dataset. Up until now, the antibiotic dose was administered in full at the beginning of the experiment. This type of experiment will be further referred to as *single treatments*. In new experiments, we again administered a range of antibiotic concentrations at the beginning of experiment (first dose), and added a second dose at some specific time later (generally 2, 4, or 6 hours from the beginning). These experiments will be referred to as *repeated treatments*.

As new calibration dataset, we have chosen OD data of single treatment experiments together with 3 repeated treatment experiments. To investigate the robustness of predictions obtained with this new calibration dataset, we fitted the model forty times on this new dataset and calculated the corresponding CFU prediction scores. As shown on Figure 3b (green histogram), this method allowed the optimizer to reach parameter sets with good predictive power much more consistently. These results apply to four other strains (two other strains on Figure 3c, all five tested strains in Supplementary Text 7).

What makes repeated treatment experiments so informative that they allow the model to produce good CFU predictions? One of the reasons is that they provide information about β-lactamase efficiency: with a high initial antibiotics dose, the second dose does not have much impact on time of regrowth, which shows that the fraction of dead population mainly depends on the initial dose, and that β-lactamases released by dead cells is sufficient to degrade the two doses of antibiotic. In Supplementary Text 9, we investigated the impact of the choice of the calibration dataset by comparing results based on different combinations of repeated treatment experiments in terms of fitting success, CFU prediction quality, and tightness of the estimation for key parameter values. From these results, we conclude that calibration datasets containing 3 or 4 repeated treatment experiments are a good compromise between the complexity of generating the training dataset, and the quality of the prediction score and the identifiability of model parameters.

In summary, with our slightly extended experimental approach, we have demonstrated that one can robustly calibrate our model on OD data only so that it can then predict the temporal evolution of the number of live cells during antibiotic treatments. This is a major result that has never been achieved before.

### Predicting complex temporal evolution of CFU for delayed treatments

In the previous section, we focused on predicting the temporal evolution of CFUs in single treatment experiments. As an additional challenge for the predictive power of the model, we sought to predict the temporal evolution of CFUs in delayed treatments, i.e. in treatments where the whole antibiotic dose is administered several hours after the beginning of the experiment. This is a way to test the capacity of the model to capture the inoculum effect since the bacterial population is at higher densities at the time of antibiotic administration.

On Figure 4a and in Supplementary Text 10, we show the temporal evolution of OD and CFUs for a clinical isolate treated at 0, 2, 4, or 6 hours with 16mg/L or 32mg/L of cefotaxime. When the treatment is applied immediately or after 2hrs, we observe an exponential increase of OD values for about four or five hours, then a plateau and finally regrowth up to maximal carrying capacity. Interestingly, for later treatments, the OD shows exactly the same profile as non-treated populations: continuous exponential growth up to reaching the maximal carrying capacity. Yet, the antibiotic has an effect on cells, as evidenced by the decrease in CFU counts observed for the treatments at 4 and 6 hours. At 24 hours, the OD of the cell population that was treated late is the same as the OD of non-treated cell populations, but not the CFU counts. This strongly suggests that late treated cells are growth arrested in a filamentation state and that nutrients have been exhausted before antibiotic has been degraded. Using our model calibrated as explained in the previous section, that is, calibrated on OD data only (Fig 3b, bottom), we obtain a very good agreement between predictions and experimental data on both delayed treatments (Figure 4a and Supplementary Text 10). Strikingly, the model is able to reveal that late antibiotic treatments do affect cell growth, despite the fact that OD profiles are perfectly identical as those of untreated populations.

**Figure 4.**
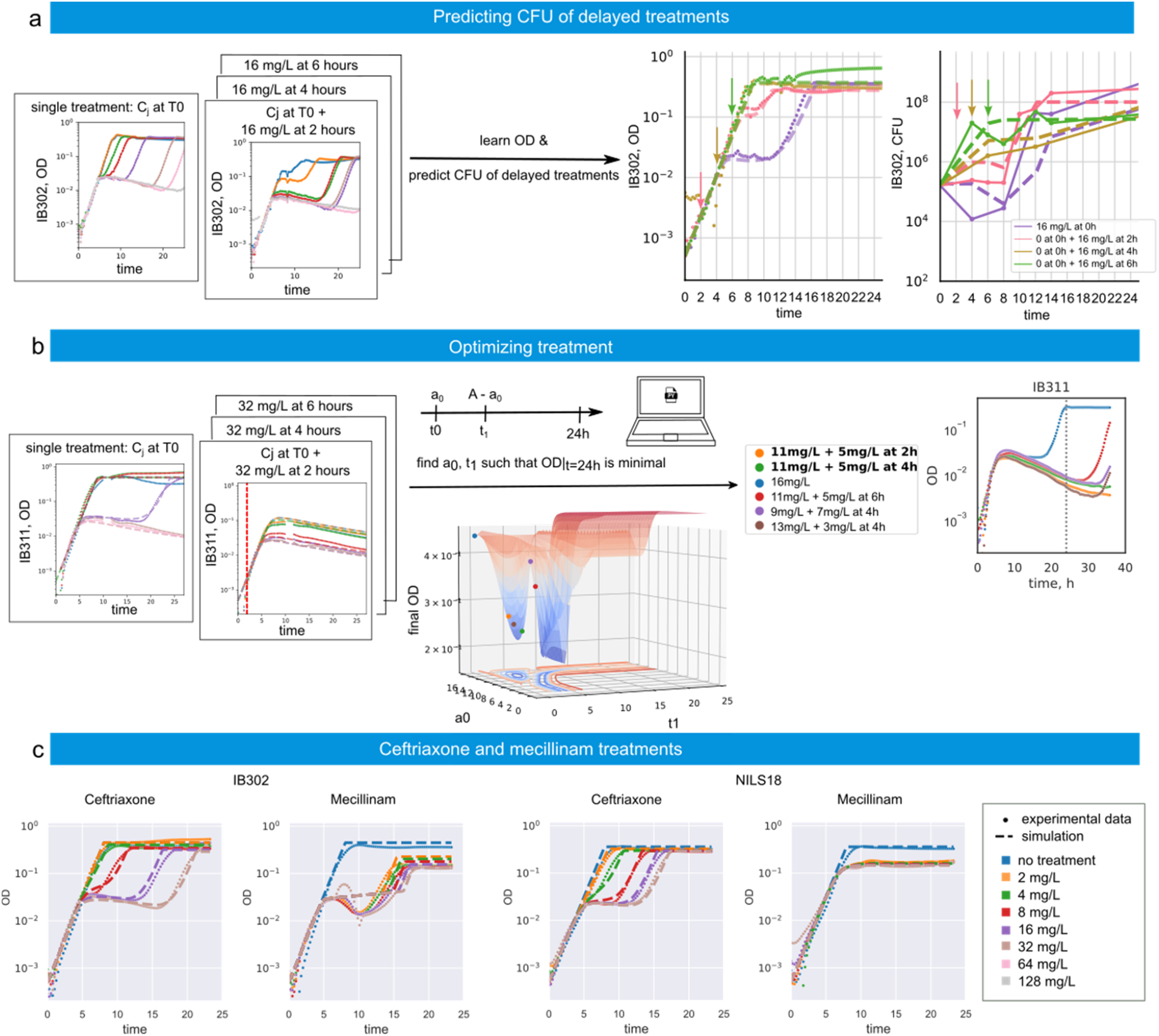
Challenging further the predictive capacities of the model. **a** Predicting complex temporal evolution of CFU for delayed treatments. The calibration dataset is represented on the left side. Experiments corresponding to the data on the plots on the right side are delayed treatment experiments, where no antibiotics are added at the beginning of experiment, and the whole amount (the same for all presented plots) is administered several hours later. Experimental data is presented in points with solid lines and simulations are represented by dashed lines. **b** Optimal treatments. We consider the following optimization problem: using one of the parameter value sets fitted on one single and three repeated treatment experiments, find a way to split a given antibiotic amount into two separate doses, one (*a*_0_) applied at the beginning and the rest applied at time *t*_1_ so that the OD at 24 hours is minimal. *Left*: calibration dataset and fitting results. *Center:* surface of the optimization search. *Right:* Experimental results. The color of lines corresponds to specific treatments listed in the table on the left. The positions of these treatments on the optimization surface are shown by the points of corresponding color. **c** For two clinical isolates (IB302 – left, NILS18 – right), we measured the temporal evolution of OD in response to treatments with two other β-lactams (ceftriaxone and mecillinam), fitted the model to the data, and presented simulated behavior. Experimental data is in points, simulations of the model is in dashed lines.

### Optimal treatments

Another challenge, that we investigated with our model, was to find optimal treatments. More precisely, we wondered how to best apply a fixed antibiotic amount (A) into two doses: one (*a*_0_) is to be applied at the beginning, the rest is to be applied later (at *t*_l_ hours), in a way that minimizes the OD at a specific time (24 or 30 hours). Actually, this problem is quite challenging since, as shown in the previous section and because of the inoculum effect, late additions of antibiotics have lesser effects on OD. Therefore, one should expect -and this is indeed generally true-that the best solution is the trivial one: applying all the dose at the initial time. Interestingly, we predicted for the IB311 strain and a 16mg/L dose, that the optimal treatment that minimizes OD at 24hrs was non trivial: 11mg/L of cefotaxime should be added at T0, and 5mg/L should be added between T2 and T4. We tested experimentally the predicted optimal solution, as well as a few different treatments selected in the neighborhood of the predicted optimal one. We found that the predicted optimal treatment was indeed the best performing of all the treatments tried, and that the response to all treatments followed model predictions (Figure 4b). Incidentally, this search for the treatment that gives the lowest OD at 24 hours seems to correlate with the treatment that delays cell regrowth the most: all tested combinations prevented regrowth before 24 hours, and the optimal solution even prevented regrowth at 30 hours. Interestingly, this is the only strain out of the five clinical isolates tested for which we managed to find a non-trivial optimal treatment. What distinguishes it from the others? Looking in details at the temporal evolution of the OD in the presence of different concentrations of cefotaxime, one can observe that this strain has some specific features: at low concentrations (up to 8 mg/L), there is no impact of antibiotic treatment on the growth dynamics, at one specific concentration in the tested range (16 mg/L), there is a crash, growth arrest and regrowth, and at higher concentrations (above 32 mg/L), there is a crash and growth arrest, but no regrowth (Figure 4b, left panel). All other tested strains show regrowth for single treatments (Figure 3c, left panels). Looking into the genotype, this strain expresses three β-lactamases (TEM-1, OXA-181, CMY-2), but does not express a β-lactamase of CTX-M type, contrary to the other four tested strains. This strongly suggests that the 3 β-lactamases are sufficient to confer a protection to the cell (single-cell resistance, periplasmic action) but not to provide a good protection to the cell population (collective antibiotic tolerance, environmental action). A more detailed analysis of strain phenotypes and model parameter values is provided in Supplementary Text 6.

When extending the time horizon, and considering optimal treatments that minimize OD at 30 hrs, our model predictions yielded another non-trivial solution for the NILS18 strain treated with 64mg/L of cefotaxime. On this time horizon, regrowth cannot be prevented. Instead, the strategy amounted to hit the population so as to kill a the highest number of cells such that the glucose used by these cells will be missing at the regrowth time and the cell population will not be able to reach the carrying capacity anymore. With strains that release β-lactamases in the environment, finding the treatment time that kills the highest number of cells is very non-trivial. In Supplementary Text 11, we provide evidence that this strategy is indeed effective.

### Ceftriaxone and mecillinam treatments

In all previously described experiments, we studied the bacterial responses to treatments with one specific β-lactam, cefotaxime, and tested the capacity of our model to capture them. To further test the generality of our model, we test treatments with two other antibiotics: ceftriaxone, a broad-spectrum cephalosporin, inhibiting cell wall synthesis and inducing filamentation; and mecillinam, an extended-spectrum penicillin, targeting primarily PBP2 and inducing production of osmotically-stable round cells^14^.

The growth kinetics assays show quite different response for the two clinical isolates to antibiotic treatments, especially for mecillinam. In the case of IB302, there is a (now classical) period of exponential growth followed by growth arrest. Biomass degradation is much more noticeable however, and time of regrowth is not very dependent on antibiotic concentration. Interestingly, treated cell populations do not regrow to full carrying capacity. In the case of NILS18, the exponential growth period is much longer than with other antibiotics and followed by growth arrest at an OD lower than the carrying capacity with no regrowth. Altogether, these dynamics are very different from the ones we saw with cefotaxime. Using growth parameters from parameter value sets previously generated on cefotaxime, we calibrated the model to new data. Simulations are in perfect agreement with experimental data corresponding to ceftriaxone treatments, which is quite expected given that ceftriaxone is very close by its mode of action to cefotaxime. However, the agreement between simulations and experiments is less perfect in the case of mecillinam, which is understandable given that mecillinam is quite different from the other two antibiotics and does not induce filamentation. Still, the model accurately captures the decrease in final OD, providing indirect evidence that the magnitude of cell death was appropriately captured too.

## Discussion

After decades of research on antimicrobial resistance, the modes of action of antibiotics and the resistance mechanisms of bacteria are now known in great detail. Surprisingly, however, this accumulated knowledge has not yet proven effective enough to predict the bacterial response to antibiotic treatments at the cellular and at the population levels with quantitative accuracy for ESBL pathogenic bacteria. This lack of accuracy necessarily limits our capacity to develop rational treatments and precision medicine approaches. Here, we proposed a model-based approach to characterize in a quantitative manner the dynamical response of ESBL *E. coli* clinical isolates to β-lactam treatments. The model, based on a growth-fragmentation equation and on a limited number of simple biological hypotheses, represents a comprehensive understanding of phenomena happening at the molecular, cellular, and population levels, as well as their interactions. Notably, it gives a central role to cellular filamentation as a way for cells to gain time until degradation of the antibiotic by the β-lactamases released in the environment by the fraction of dead cells. Our model is the first model that provides a unified explanation to observations made at the population level (OD profiles) and at the cellular level (CFU profiles). Moreover, it has a strong predictive power. When calibrated using a slight extension of OD-based data that we propose here, it can predict the CFU profiles in initial and delayed treatments despite the inoculum effect, and suggest non-trivial optimal treatments. The capacity to predict CFUs from OD measurements could have important practical implications given the difficulty of performing CFU assays in comparison to growth kinetics assays.

This work addresses an observational and modelling challenge never tackled head-on by the community. Past quantitative works focused on specific aspects of bacterial response to β-lactams but do not have our integrative viewpoint and strong predictive power. Lee and colleagues focused on the quantitative characterization of growth and death rates of cells subjected to β-lactams and found a linear correlation between these two rates for susceptible cells (lab strains or ESBL strains treated in presence of β-lactamase inhibitors)^10^. A model is proposed to support the intuition that this observed relation is consistent with a mechanism by which the antibiotic causes filamentation and filamentation increases cell lysis probability, a mechanistic explanation that we use here as well. The model is not fitted to actual data. Going further, Kim and colleagues characterized the death probability as a function of the cell length using videomicroscopy data^11^. A model is used semi-quantitatively to predict the temporal evolution of the live biomass for susceptible cells. The model is not fitted to temporal data. It would lack key processes, notably antibiotic degradation by released β-lactamases. And precisely, Meredith and colleagues focused on quantifying the relative importance of single-cell resistance and collective antibiotic tolerance effects for population survival^16^. Again, a model is provided to support the intuition. It is not calibrated to data and lacks key features like cell filamentation and the masking of the live biomass dynamics by the remaining dead biomass. Lastly, Baig and colleagues proposed a neural network approach to capture key features in microbial growth dynamics, and among other applications, use it to as a model fitting tool for the calibration of an ODE model of cell population response to strong amoxicillin treatments^17^. The model allows to capture qualitatively the resistant or susceptible profile of the tested isolates. Its quantitative accuracy is harder to establish, however, the information being provided on the difference of the derivative of the OD. The average relative error on the derivative of the OD is 14.5% at all time points. This work can nevertheless inspire us to replace the complex ODE approximations of the initial PDE model by a neural network alternative model using physics-informed machine learning to form an hybrid neural network^18,19^. Such an approach could further increase the accuracy of our model while preserving the interpretability of the parameters appearing in the mechanistic part.

Our model captures in an elaborate manner the cell response to complex treatments (pharmacodynamics) but also the evolution of the cells’ environment, that is, of the glucose, of the antibiotic and of the β-lactamases concentrations in the culture medium in the well of a microplate (elementary pharmacokinetics). By completing our model with a proper pharmacokinetics component, we would obtain an excellent starting point to quantitatively investigate the effects of treatments *in vivo*, in which our model could notably help test whether the local release of β-lactamases at the site of infection can have a real impact on treatment outcomes. There is indeed a known correlation between the presence of an *in vitro* inoculum effect and failures of *in vivo* treatments^20^.

## Methods

### Strains and antibiotics

The 11 *E. coli* clinical isolates used in this study have different origins (feces, urine, blood) and different genetic backgrounds. These strains express between 1 and 5 β-lactamases. They are described in detail in Supplementary Text 1. Cefotaxime was purchased from Sigma-Aldrich and dilutions were made considering the purity specified by the vendor. Antibiotics and stock solutions were kept at -20°C to limit degradation.

### Growth conditions

Unless specified otherwise, all pre-cultures and cultures were performed in 0.1% glucose M9 liquid medium (1 g/L glucose, 6.78 g/L Na_2_HPO_4_, 3 g/L KH_2_PO_4_, 1 g/L NH_4_Cl, 0.5 g/L NaCl, 0.24 g/L MgSO_4_, 0.01 g/L CaCl_2_), and experiments were performed in 0.1% glucose M9 liquid medium with 0.1% Tween 20. The low glucose concentration is used to create the conditions of a carbon-related growth arrest, a situation that lends itself better to mathematical modelling (Monod growth). The growth arrest happens in this medium between 0.2 and 0.3 OD_600_. For overnights and precultures, cells were incubated at 37°C and at 200 rpm.

### Growth curves acquisition

**Overnight**. A single bacterial colony was picked from an agar plate and incubated overnight at 37°C and 200 rpm. **Preculture**. The overnight was vortexed and diluted to 0.05 OD_600_ before a new incubation of 3 hours under the same conditions, aiming to obtain the cells in exponential phase at the beginning of the experiment. **Experiment preparation**. Cells from the preculture were vortexed and diluted to 0.01 OD_600_. Each well of a 96-well plate with transparent flat bottoms was filled with 190 μL of antibiotic dilution in M9 media with Tween 20 and 10 μL of the cell suspension. Depending on the plate configuration and the number of replicates, the dilutions were made as much as possible in larger quantities in order to minimize errors related to small-volume pipetting. **Data collection**. Tecan Spark multimode plate readers were used for all OD acquisitions. Thermo Scientific™ Nunc™ Edge 2.0 flat bottom microtiters plates were used. OD measurements were carried out in a loop consisting of measurement and incubation with shaking. The incubation periods are carried out at 37°C and last 300 s. **Repeated treatments**. Stocks of antibiotics were prepared in advance and kept at +4°C. Before application they were warmed up at room temperature, then put into a small container. From there, using an Integra Biosciences Voyager 8-channel electronic pipette, antibiotic was administered to a column of the 96-well plate at the same time to limit the time outside of incubation conditions.

### Live cell number estimation

Live cell numbers have been estimated by the CFU counting method. At every timepoint the plate was removed from the plate reader, 5 μl were sampled using an Integra Biosciences MINI 96 electronic pipette with 96 channels into 45 μl of PBS buffer. The experimental plate was put back into the plate reader and the experimental program was resumed. The previous dilution was mixed. 5 μl were spotted in a square dish filled with Lysogeny broth (LB) agar. 10 μl were transferred into 90 μl of PBS in the next plate, mixed, and 5 μl were spotted on agar plates. This step is repeated to obtain 6 different dilutions. All agar plates were kept at room temperature until the last timepoint, when all the plates were put for incubation at 37°C for the night. The next day the plates were photographed on a black background. The resulting pictures have been analyzed with CFU Spot Reader, an Ilastik and R-Shiny-based application that extracts colony counts from photographs of droplet-based CFU experiments. Relevant colony counts have been manually validated. CFU Spot Reader has been developed by Vincent Aranzana-Climent and the U1070 Pharmacology of Antimicrobial Agents and Antibioresistance team at INSERM. The code is available upon request.

### Data analysis

**OD blanking**. The measured OD of a well is the sum of the OD of the cell culture, and of the OD of the well bottom, which slightly varies among plates. For this reason, before an experiment, the OD of a plate filled with media is measured and a constant OD corresponding to an average OD of the plate without bacteria was removed from each well data. **Removal of aberrant timepoints**. Occasionally, an OD reading fails and returns a value much larger than both the previous and the following ones of the same well. As this situation is highly biologically improbable, these outliers were removed for model fitting steps.

### Model

In the model, we denote by *s, a, b, c*, and *c*_r_, the concentrations of nutrients, antibiotics, and β-lactamases, and the dead degradable and non-degradable biomasses, respectively. We also denote by *n*(*l, t*) the extensive cell length distribution at time *t*. The total number of cells is then 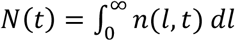, and the total live biomass is 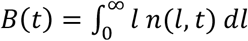. The dynamics of the first five state variables are described by the following ordinary differential equations:

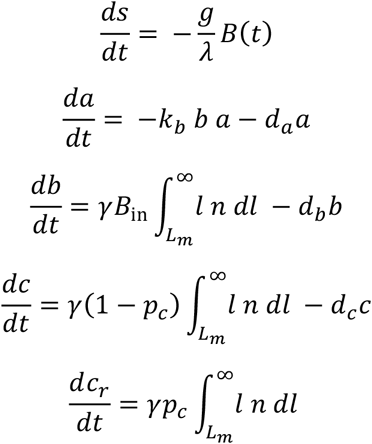

with the growth rate *g*, following Monod equation, and the optical density *OD*, assumed proportional to the sum of the live and dead biomasses, defined as follows:

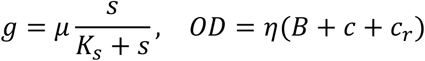

To describe the temporal evolution of the distribution of cell lengths, *n*(*l, t*), we used a growth-fragmentation model. It relies on four main mechanisms: the cells continuously elongate with rate *g*, they divide with rate *f* and when they divide, regardless of their length, they always split into cells of sizes between 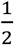 and 1 (arbitrary unit), meaning that filamented cells might split into more than 2 cells. Finally, they lyse when they reach the critical length *L*_*m*_. The following PDE describes these phenomena:

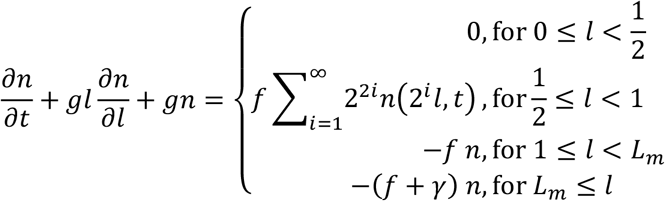

The PDE is initialized with the steady state cell length distribution of an exponentially growing cell population: *n*(*l*, 0) = *N*(0) *y*_∞,*γ*=0_(*l*) with

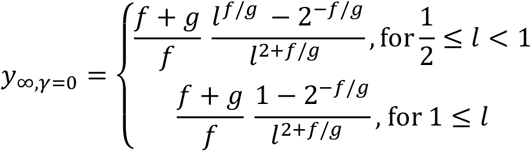

The interaction with the antibiotics is taken into account through the dependency of these rates and threshold with the antibiotics concentration in the culture medium. While the elongation rate is independent on the antibiotic concentration and simply follows Monod equation, the division rate *f* and the critical length *L*_*m*_ are decreasing functions of the antibiotic:

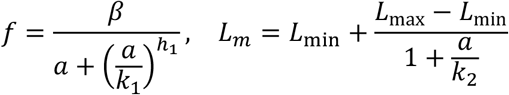

This describes the complete PDE model. For computational efficiency, we derived and used for calibration an ODE model approximating the PDE model above (see Supplementary Text 2 for detailed explanations).

### Model fitting

Each parameter was restricted to a range of biologically plausible values. Depending on the role of the parameter and the size of the range, a change of variable was applied or not, to perform the search of this parameter in the linear space or in a logarithmic space. All 17 search ranges were then brought back to the interval [0,10] and it is in this [0,10]^l7^ space that the parameter search was performed.

The cost function computes the log-likelihood of the data assuming independent Gaussian measurement noise on each point. Special care was taken when searching on mixed datasets involving both OD measurements and cell counts, to balance the contribution of both data types. The integration of the ODE system was done with the ‘diffeqsolve’ method of highly-efficient diffrax package with the method ‘Tsit5’, and absolute and relative tolerances set to 10^-6^ (Ref^21^).

Model fitting was performed both with CMAES^22^, with an initial *σ* = 2, and Latin Hyper Cube with 5000 samples and method ‘trf’. More specifically, model calibration was done in two steps. First, out of the complete set of model parameters, we selected a subset that is responsible for normal growth of a bacterial population without antibiotic present (maximal growth rate *μ*, maximal division rate *β*, half-velocity constant of nutrients *K*_*s*_, conversion factor from nutrients *λ* and conversion rate between OD and CFU *η*). The other parameters being irrelevant to describe the growth kinetics of bacteria without antibiotics, we arbitrarily fixed their values and calibrated this selected subset of parameters to OD and CFU data of a growth kinetics without cefotaxime present. Secondly, we relaunched the search for optimal parameter values on OD and CFU data fixing the growth parameters to the ones found during the first step.

For model prediction and treatment optimization purpose, we performed the fits 50 times using the OD data of one single and three repeated treatment experiments (as in Figure 3c), removed from the resulting parameter sets a few sets exhibiting unrealistic OD profiles, and selected parameter sets having the best prediction score (lowest CFU cost) or the best combined score (lowest sum of OD fitting cost and of CFU prediction cost).

## Data availability

All experimental data generated in this study have been deposited on Zenodo: https://doi.org/10.5281/zenodo.12543534.

## Code availability

The Python code to process and analyze the raw data and generate the figures of the manuscript can be found at https://gitlab.inria.fr/InBio/Public/esbl-escape. It uses numpy, scipy, pymc, matplotlib, pandas, mpmath, cma, pyyaml, jax, jaxlib, diffrax, ipython, equinox, openpyxl, pyDOE, seaborn, and scikit-learn packages. The code for CFU Spot Reader is available upon request at.

## Acknowledgements

The authors would like to thank Erick Denamur for discussions and clinical isolates. They also thank Vincent Aranzana-Climent, Nicolas Grégoire and Julien Buyck for giving us an early access to their code, and for their help and guidance with installation and adaptation of the tool. This work was supported by ANR grants Inception (ANR-16-CONV-0005), Anoruti (ANR-20-PAMR-0001), and Seq2Diag (ANR-20-PAMR-0010).

## Author information

These authors contributed equally: Virgile Andreani and Viktoriia Gross; Imane El Meouche and Gregory Batt.

Author contributions: VA, VG, IEM and GB designed the research. VA developed the PDE model and derived the ODE approximation. VA performed a preliminary set of experiments and their analysis. VG developed protocol improvements. VG performed all the experiments presented in the manuscript and their analysis. VA and VG developed code for data analysis, efficient model calibration, and calibration result analysis. LY discussed the project and provided expertise on escape mechanisms. PG advised the project, provided expertise on b-lactam resistance and for the choice of the isolates. IEM contributed to protocol design and to critical interpretation of the results. IEM and GB supervised the research. VG and GB wrote the article, based on an initial version made by VA and GB, and with significant input from all other authors.

## Ethics declarations

The authors declare no competing interests.

