## Supplementary Information for "A quantitative approach to easily characterize and predict cell death and escape to β-lactam treatments"

**Virgile Andreani[1*§], Viktoriia Gross[1,2*], Lingchong You[3], Philippe Glaser[4], Imane El Meouche[2#], Gregory Batt[1#]**

1. Institut Pasteur, Inria, Université Paris Cité, Paris, France
2. Université Paris Cité, Université Sorbonne Paris Nord, Inserm, IAME, Paris, France
3. Department of Biomedical Engineering and Center for Genomic and Computational Biology, Duke University, Durham, NC, USA
4. Unité EERA, CNRS UMR 3525, Institut Pasteur, APHP, Université Paris–Saclay, 28 rue du Docteur Roux, 75015 Paris, France

* and # These authors contributed to this work equally

§Present address: Biomedical Engineering Department, Boston University, Boston, USA

**Supplementary Text 1: Detailed information on *E. coli* clinical isolates**

The strains used in this work have been sequenced as a part of previous studies: Ref[1] for IB strains and Ref[2] for NILS strains. Antibiotic-related genes and mutations are known. Table 1.1 represents a summary of available information for each strain.

| Strain | Origin | ST Warwick | β-lactamases and Carbapenemases | Other relevant antibiotic resistance genes | Mutations contributing to susceptibility |
|---|---|---|---|---|---|
| NILS1 | Feces | 69 | TEM-1B | aph6-Id, aph3"-Ib, aadA5, mphA, mdfA, catA1, sul1, sul2, tetB, dfrA17 | |
| NILS11 | Feces | 10 | TEM-1B | aph3"-Ib, mdfA, sul2, dfrA5, dfrA1 | |
| NILS12 | Feces | 450 | TEM-1B | aph6-Id, aph3"-Ib, aph3'-Ia, aac3-Iid, aadA5, mdfA, mphA, sul1, sul2, tetA, dfrA17 | gyrA S83L/D87N, parC S80I |
| NILS18 | Blood | 69 | CTX-M-14, TEM-1B | aph6-Id, aph3"-Ib, mdfA, mphA, sul2, tetB, dfrA14 | gyrA S83L, parC S80I |
| NILS42 | Urine | 38 | CTX-M-14b, TEM-1B | aph6-Id, aph3"-Ib, aadA1, mdfA, sul2, dfrA1, dfrA5 | |
| NILS56 | Urine | 405 | CTX-M-15 | mdfA, tetA, tetB | gyrA S83L/D87N, parE S458A, parC S80I |
| NILS64 | Urine | 131 | CTX-M-15, TEM-1B, OXA-1 | aac6'-Ib-cr, aac3-IIa, aadA5, mdfA, mphA, catB3, sul1, tetA, dfrA17 | gyrA S83L/D87N, parE I529L, parC S80I/E84V |
| IB302 | Unknown | 410 | TEM-1B, CTX-M-15, OXA-1 | aac(6')Ib-cr; dfrA17; sul2; aadA5; aac(3)-IId; strA; sul1; strB; tet(B); mph(A) | gyrA S83L/D87N, parC S80I |
| IB307 | Rectal Swap | 410 | TEM-1B, CTX-M-15, OXA-1, NDM-5, CMY-42 | aadA2; dfrA17; aac(6')Ib-cr; tet(A); erm(B); strA; tet(B); sul1; sul2; aadA5; strB; mph(A); qepA; dfrA12 | ftsI YRIK, gyrA S83L/D87N, parC S80I/E84 |
| IB308 | Rectal Swap | 410 | CTX-M-15, OXA-181, CMY-42 | QnrS1; sul2; mph(A); tet(A); sul1; aadA5; dfrA17 | ftsI YRIK, gyrA S83L/D87N, parC S80I/E84 |
| IB311 | Rectal Swap | 410 | TEM-1, OXA-181, CMY-2 | QnrS1; tet(B); dfrA17; mph(A); sul2; aadA5; aac(3)-IId; strA; sul1; strB | ftsI YRIN_349-532, ompC R195L, ompF -46; C->T (OmpR F3), gyrA S83L/D87N, parC S80I/E84 |

**Table 1.1** Main features of the clinical isolates used in this study.

The gene coding for PBP3 is *ftsI*. A mutation in this gene can lead to reduced susceptibility of PBP3 to β-lactams. *ompC* and *ompF* respectively code for a precursor of the outer membrane porins C and F. Their mutations can play a role on antibiotic susceptibility because they might prevent the entrance of antibiotic molecules in the cell. *gyrA* and *parC* are genes coding for a DNA gyrase and topoisomerase, enzymes that participate in the winding and unwinding of DNA. These mutations are not involved in the resistance to β-lactams, but rather to fluoroquinolones.

**Supplementary Text 2: Mathematical models of bacterial population response to β-lactam treatments**

The models have been built on several core hypotheses, motivated by literature and experimental evidence, developed in the main text and summarized here.

- Growth: the elongation speed per cell is proportional to the cell length (exponential growth of biomass), such that the temporal derivative of the size increase of a cell of length $l$ is $gl$. The growth rate $g$ depends on nutrients but is *not affected* by the β-lactam antibiotic for the antibiotics considered[3,4].
- Division: β-lactams, through their action on PBP3, affect the ability of the cells to divide[5]. Therefore, the never stopping biomass formation leads to cell filamentation if the antibiotic concentration is large enough[6]. In the model, the division rate is represented by the function $f$, which is a decreasing Hill function of the antibiotic concentration $a$.

- Division of filamented cells: as confirmed in Ref[7], divisions of filamented cells after antibiotic removal happen faster than for shorter cells, and split the cell into sizes that correspond to "units" of normal cells. This process lends itself well to modelling if we consider that cells of sizes between $2^{i-1}$ and $2^i$ divide into $2^i$ equal cells, as shown in Supplementary Figure 2.1.
- Death: The death rate of cells depends on their capacity to maintain the integrity of their cell wall and on their actual length. More precisely, we modelled this phenomenon by a constant death rate that cells experience above a critical length $L_m$. This critical length decreases when antibiotic concentrations increase, which we interpreted as the disruption of the action of the wall-repairing enzyme PBP1. Interference with the action of PBP1 is indeed known to result in rapid cell lysis[6].
- Collective antibiotic tolerance: Bacteria release β-lactamases in the environment upon lysis[8]. These enzymes are able to degrade the antibiotic molecules in the cell culture[9], thus protecting the bacteria not lysed yet.

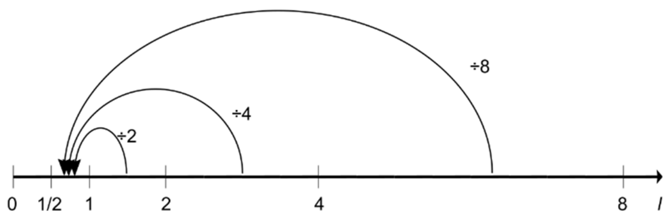

**Figure 2.1** Schematic representation of the division of cells in the model.

It stems from these hypotheses that the model needs to account for cell lengths and their heterogeneity in the cell population. Therefore, we developed a growth-fragmentation model, expanding on the preliminary work of Hall and Wake[10]. The full PDE model is as follows:

$$\frac{\partial n}{\partial t} + gl\frac{\partial n}{\partial l} + gn = \begin{cases} 0, & \text{for } 0 \le l < \frac{1}{2} \\ f\sum_{i=1}^{\infty} 2^{2i} n(2^i l, t), & \text{for } \frac{1}{2} \le l < 1 \\ -f\, n, & \text{for } 1 \le l < L_m \\ -(f+\gamma)\, n, & \text{for } L_m \le l \end{cases}$$

$$\frac{ds}{dt} = -\frac{g}{\lambda}\int_0^\infty l\, n\, dl$$

$$\frac{da}{dt} = -k_b\, b\, a - d_a a$$

$$\frac{db}{dt} = \gamma B_{in}\int_{L_m}^\infty l\, n\, dl\ - d_b b$$

$$\frac{dc}{dt} = \gamma(1-p_c)\int_{L_m}^\infty l\, n\, dl\ - d_c c$$

$$\frac{dc_r}{dt} = \gamma p_c \int_{L_m}^\infty l\, n\, dl$$

with

$$g = \mu\frac{s}{K_s + s}$$

$$f = \frac{\beta}{a + \left(\frac{a}{k_1}\right)^{h_1}}$$

$$L_m = L_{\min} + \frac{L_{\max} - L_{\min}}{1 + \frac{a}{k_2}}$$

$$OD = \eta\left(\int_0^\infty l\, n\, dl + c + c_r\right)$$

where $n(l,t)$ is the extensive cell length distribution at time $t$, $s$ the concentration of nutrients in the culture medium, $a$ the concentration of β-lactams, $b$ the concentration of β-lactamases, $c$ and $c_r$ the OD of dead biomass (degradable and not degradable). The total number of cells is $N(t) = \int_0^\infty n(l,t)dl$.

Some initial conditions are given by the experiment: $s(0), a(0), N(0)$ (the initial total number of cells), and $b(0) = c(0) = c_r(0) = 0$. Because we start the experiment with cells in exponential phase, we take as initial condition for $n$ its steady size distribution with no death: $n(l,0) = N(0)\, y_{\infty,\gamma=0}(l)$ with

$$y_{\infty,\gamma=0} = \begin{cases} \frac{f+g}{f} \frac{l^{f/g} - 2^{-f/g}}{l^{2+f/g}}, & \text{for} \frac{1}{2} \leq l < 1 \\ \frac{f+g}{f} \frac{1-2^{-f/g}}{l^{2+f/g}}, & for\ 1 \leq l \end{cases}$$

A simulation of this model can be shown on Supplementary Figure 2.2.

The simulation of this PDE requires a custom numerical method. Despite taking care of its efficiency, the simulation of this equation takes a few seconds for a single initial condition, and more than a minute for 12 initial conditions, which corresponds to the number of initial conditions in a typical experiment. However, to extract parameter values from the experimental data, the model needs to be simulated a large number of times, for different parameter sets. This is difficult if the simulation is so computationally expensive. For this reason, we proceeded to simplify the model to make it easier to simulate.

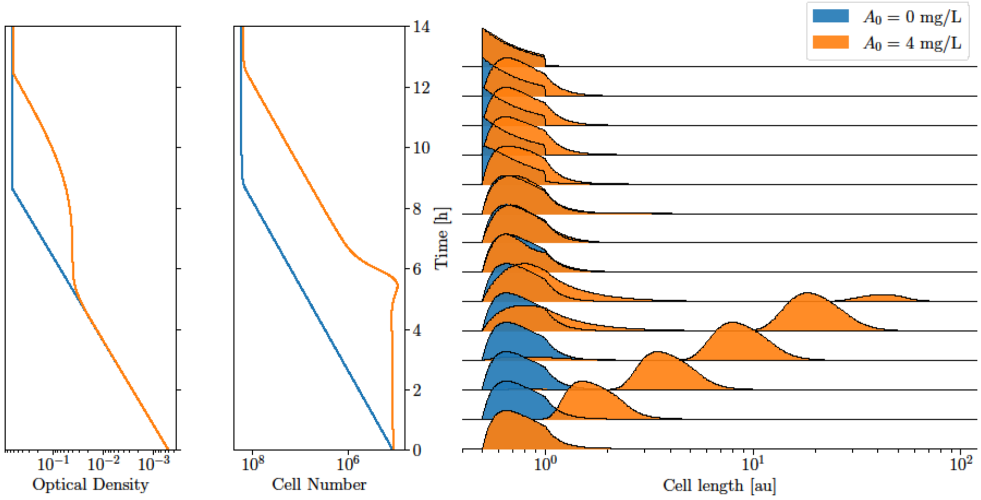

**Supplementary Figure 2.2 Simulation of the PDE model for two different initial antibiotic doses.** We plot the evolution in time of the OD, of the number of cells, and of the distribution of cell lengths. The untreated population shows exponential growth both in OD and cell number, and its length distribution remains stable until the nutrients start lacking between 8 and 9 hours. Then, it transitions to its stationary length distribution. The treated population shows an exponential growth in OD (increase in biomass), but its cell number remains constant, and its length distribution is shifted towards longer cells, during the first 5 hours. At this point, the antibiotic has been cleared and the initial distribution is rebuilt, and the cell number starts growing exponentially, which comes eventually with a raise of the OD, until the nutrients start lacking between 12 and 13 hours.

The computationally expensive part is the simulation of the entire cell length distribution, represented as a continuous function on positive real numbers. A way to transform this distribution into a more manageable object is to represent it with a finite number of variables: its first moments. We then replaced $n(l,t)$ by the total number of cells $N(t) = \int_0^\infty n(l,t)\, dl$ and their average length $L(t) = \int_0^\infty l\, n(l,t)\, dl\, / N(t)$.

This transformation gives the proper form for the feedback of death on $N$and $L$. Working through the elimination of $n(l,t)$, we arrive at a system of ODEs involving $N, L$, and two partial moments which we called $Y_>(t)$ and $L_>(t)$. The expression of these moments still depends on the length distribution $n(l,t)$. Here, with careful approximations, we managed to approximate these two partial moments by two functions of $L$ only (see Andreani for more details[11]). The resulting ODE model is the following, which is the one that we used for parameter inference.

$$\frac{dN}{dt} = N\left[f\left(\frac{L}{\ln 2} - 1\right) - \gamma Y_>\right]$$

$$\frac{dL}{dt} = L\left[g - f\left(\frac{L}{\ln 2} - 1\right)\right] - \gamma(L_> - LY_>)$$

$$\frac{ds}{dt} = -\frac{g}{\lambda} NL \qquad \frac{db}{dt} = \gamma B_{in} N L_> - d_b b$$

$$\frac{da}{dt} = -k_b b\, a - d_a a \qquad \frac{dc}{dt} = \gamma(1-p_c) N L_> - d_c c$$

$$\frac{dc_r}{dt} = \gamma p_c N L_>$$

$$g = \mu \frac{s}{K_s + s}$$

$$\nu = \frac{\beta}{g}$$

$$f = \frac{\beta}{1 + \left(\frac{a}{k_1}\right)^{h_1}}$$

$$L_m = L_{min} + \frac{L_{\max} - L_{\min}}{1 + \frac{a}{k_2}}$$

$$L_0 = \left(1 + \frac{\mu}{\beta}\right) \ln 2$$

$$OD = \eta(NL + c + c_r)$$

$$Y_>\left(x = \frac{L}{L_0 L_m}\right) = \begin{cases} \frac{x}{\nu}\left(x^\nu - \left(\frac{x}{2}\right)^\nu\right), & \text{for } x \leq 1 \\ x - 1 + \frac{x}{\nu}\left(1 - \left(\frac{x}{2}\right)^\nu\right), & \text{for } 1 \leq x \leq 2 \\ 1, & \text{for } 2 \leq x \end{cases}$$

$$\frac{L_>}{L_0 L_m}\left(x = \frac{L}{L_0 L_m}\right) = \begin{cases} \frac{x}{\nu \ln 2}\left(x^\nu - \left(\frac{x}{2}\right)^\nu\right), & \text{for } x \leq 1 \\ x\frac{\ln x}{\ln 2} + \frac{x}{\nu \ln 2}\left(1 - \left(\frac{x}{2}\right)^\nu\right), & \text{for } 1 \leq x \leq 2 \\ x, & \text{for } 2 \leq x \end{cases}$$

The variables and parameters of the model are presented in the following tables. All parameters are shared between the two models. The only difference in variables is $n(l,t)$ in the PDE model that becomes $N(t)$ and $L(t)$ in the ODE.

| Variable | Unit | Comment |
|---|---|---|
| $t$ | h | Time |
| $l$ | 1 | Cell length [au] |
| $n(l,t)$ | 1 | Population density |
| $s(t)$ | g/L | Concentration of nutrients |
| $a(t)$ | mg/L | Concentration of antibiotics |
| $b(t)$ | mg/L | Concentration of β-lactamase |
| $c(t)$ | 1 | Dead degradable biomass |
| $c_r(t)$ | 1 | Dead non-degradable biomass |

| Parameter | Unit | Comment |
|---|---|---|
| $\gamma$ | 1/h | Death rate |
| $\beta$ | 1/h | Maximal division rate |
| $\mu$ | 1/h | Maximal growth rate |
| $K_s$ | g/L | Half-velocity constant of nutrients |
| $\lambda$ | L/g | Conversion factor from nutrients |
| $k_1$ | mg/L | Concentration of antibiotics needed to stop cell division |
| $h_1$ | 1 | Hill coefficient of this antibiotic action |
| $k_2$ | mg/L | Concentration of antibiotics needed to stop defect repair |
| $B_{in}$ | mg/L | Concentration of β-lactamase released by a cell of length 1 |
| $k_b$ | L/mg/h | Activity rate of β-lactamase |
| $d_a$ | 1/h | Degradation rate of antibiotics |
| $d_b$ | 1/h | Degradation rate of β-lactamase |
| $d_c$ | 1/h | Degradation rate of dead biomass |
| $p_c$ | 1 | Proportion of non-degradable dead biomass |
| $L_{min}$ | 1 | Minimal cell length where lysis can occur |
| $L_{max}$ | 1 | Maximal viable cell length |
| $\eta$ | 1 | Conversion between biomass and OD |

**Supplementary Text 3: Preventing biofilm formation to improve measurement quality and reproducibility**

While performing our experiments in a 96-well microplate in M9 media with 0.1% glucose, we observed a lot of well-to-well variability: using the same media, same antibiotic and same bacterial inoculum in the same micro-plate, but in different wells, gave quite different results in terms of regrowth after antibiotic treatment. Considering that our approach consists in calibrating a model on experimental data and testing our model's capacity to capture the dynamics or even to predict some complex and previously unseen behavior, it is crucial to have data of good quality and high reproducibility. In order to investigate the cause of this variability, at the end of the experiment we performed a complete OD scan of the bottom of each well of the experimental plate and we discovered inhomogeneous patterns. Further staining with Crystal Violet revealed that these patterns were caused by biofilms. For Crystal Violet staining, first we emptied the experimental plate and washed it several times with PBS. Then we filled each well with 35 µl of Crystal Violet diluted to 0.1% in 10% ethanol and incubated in the dark for 10 minutes. Then, we emptied the plate again and washed 5 times with water. After an hour of drying in the dark, the plate was filled with 100 µl of 50% ethanol in each well and left for 10 minutes in the dark. Final step consisted in measuring OD at 595 nm[12].

A known method to prevent biofilms is to add Tween 20 into the media. After reviewing several publications and performing our own experimental tests, we selected a concentration of 0.1% that is commonly used in bio-reactor platform[3] and that was shown to prevent biofilm formation without affecting growth rate[4]. We repeated the same experiments as before, but this time with Tween 20 in the media. Well-to-well variability was significantly reduced, OD scans of well bottoms at the end of experiment showed no patterns, and, finally, Crystal Violet staining confirmed the absence of biofilms.

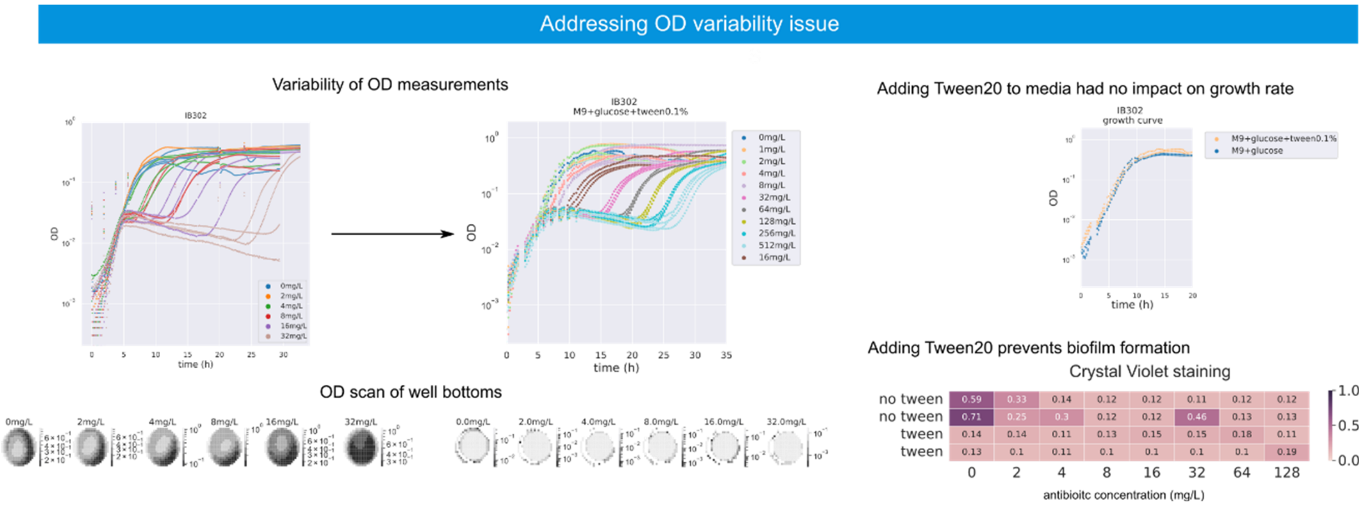

**Supplementary Figure 3.1 Addressing OD variability issue through application of Tween 20.** *Well-to-well variability of OD measurements.* Left: Characterization of bacterial response to cefotaxime treatments, different colors correspond to different antibiotic concentrations, same colors correspond to same experimental conditions. Right: Same experiment, but with addition of 0.1% of Tween 20 to the media. *OD scans of well bottoms:* for each well, OD was measured in 308 different positions; dense bacterial populations are indicated by light colors. *Impact of Tween 20 on the growth rate:* We compared growth curves in absence of antibiotic with and without Tween in the media and concluded that there is no impact on growth rate. *Crystal Violet staining for biofilm detection:* The first and last two rows correspond to experiment without and with Tween20, respectively. Different columns correspond to different concentrations of antibiotic. Values correspond to OD at 595 nm of the final step of Crystal Violet staining.

**Supplementary Text 4: Protocol for efficient droplet-based CFU assays**

One experimental plate contains 2 technical replicates of 48 different conditions (for example 6 different strains subjected to 8 different concentrations of antibiotic). In order to see the temporal evolution of the number of live cells, we need to make CFU assays at different timepoints. In our experimental protocol, we are sampling every 2 hours for the first 12 hours. In the same plate, depending on the concentration we can have biomass close to full grown stationary phase ($10^9$ CFU/mL) and almost fully lysed population ($10^3$ CFU/mL), this forces us to do up to 6 serial dilutions to capture the dynamics for all antibiotic concentrations. Therefore, using standard methods and using 1 agar-filled Petri dish per dilution per condition, for one experiment, we would obtain 96 conditions x 6 timepoints x 6 dilutions = 3456 Petri dishes to plate and count. This motivated us to pass to a droplet-based protocol.

Our experimental protocol consists in several steps: sampling, diluting and spotting on agar-filled plates, for all these steps, we are using a 96-channel electronic pipette (Integra Biosciences MINI96). First, we take a 5 µL sample from the experimental plate and dilute it in 45 µL of PBS in another 96-well plate. Before starting the next step, tips are washed with ethanol and PBS (ethanol, then air-dry, then PBS). We added this intermediate step to reduce plastic consumption and reduce experiment cost. After the washing step, we mix the last-performed dilution, then transfer 5 µL on agar-filled plate to form drops (1 drop per experimental condition), and, finally, 10 µL of this culture is diluted in 90 µL of PBS for the next dilution. Afterwards, the tips are washed again. These steps are repeated to obtain six tenfold dilutions. All prepared agar plates stay at room temperature until the last timepoint, and then plates are placed into incubator at 37°C for overnight incubation. Our goal is to limit the variations between colony sizes from different timepoints due to difference in time spent after sampling, to ease droplet analysis and automated colony counting. The next day all the plates are photographed, bottom up, on a black background.

In order to count the grown colonies and, therefore, measure CFUs, we employed CFU Spot Reader. This Shiny-based application has been developed by Vincent Aranzana-Climent and the "Pharmacology of Antimicrobial Agents and Antibioresistance" team in INSERM at Poitiers (France). This software allows to obtain CFU/mL values for each experimental condition from raw images of agar plates, corresponding to different dilutions and different timepoints, and a table specifying the experimental conditions for each position on the plate. First, each image passes through ImageJ Fiji[13] macros to be cropped, resized, and transformed from black and white into color. Second, the image goes through pixel classification in ilastik[14], a user-friendly GUI for machine learning algorithms, that separates pixels into 3 classes: background, core or edge of bacterial colony. Next, raw image together with probability map from pixel classification goes through object classification also using ilastik. Here, objects formed by pixel classified as core of a colony, are labelled as 1 bacterial colony, 2 or 3 colonies grown together, an uncountable number of colonies stuck together or an unknown object that is not a colony. From this step, we obtain for each image a list of detected objects together with information about their labelled class, their position and their size. Afterwards, an R script divides images into 96 sectors assigning a coordinate to each sector, counts colonies corresponding to detected in that sector objects, assembles information about the dilution and experimental conditions from filename and descriptive table provided at the input, and displays results through graphical interface for a user to verify manually all counted results. Final counts are further analyzed using Python scripts and used for model calibration or testing model predictive capacities. For our project, we retrained both ilastik projects on our data, and added a "not-a-colony" object. In addition, we adapted the R script to our project (our filename format, our type of experimental conditions and corresponding data, our number of sectors and our way to reference them). All relevant colony counts have been manually validated.

We performed experiments in order to test spotting and dilution variabilities. We took a bacterial culture at stationary phase, vortexed it and did two types of dilutions. On one hand, we filled a microplate with 100 µL of this culture per well, and then we performed six tenfold dilutions in PBS buffer using the protocol described earlier. On the other hand, we did two thousandfold dilutions in tubes and we filled a microplate with this diluted culture, 100 µL per well. In order to test the variability of the dilutions, we compared the results of these two different dilution protocols taking the second one as the ground truth. In order to test the variability of spotting, the last dilution from both protocols was spotted 3 times on separate plates. Results of this experiment are presented in Figure 2.1. By comparing the results from plate replications (results of the same color but different color nuance), we can conclude that spotting variability is quite low. By comparing the mean results of the two different dilution protocols, we can observe that, on average, both methods give the same result (97% accuracy when calculated per dilution), however, as expected, doing 6 dilutions with 96-channel pipette introduces more well-to-well variability. A comparable study has been recently performed by Aranzana-Climent and colleagues[15].

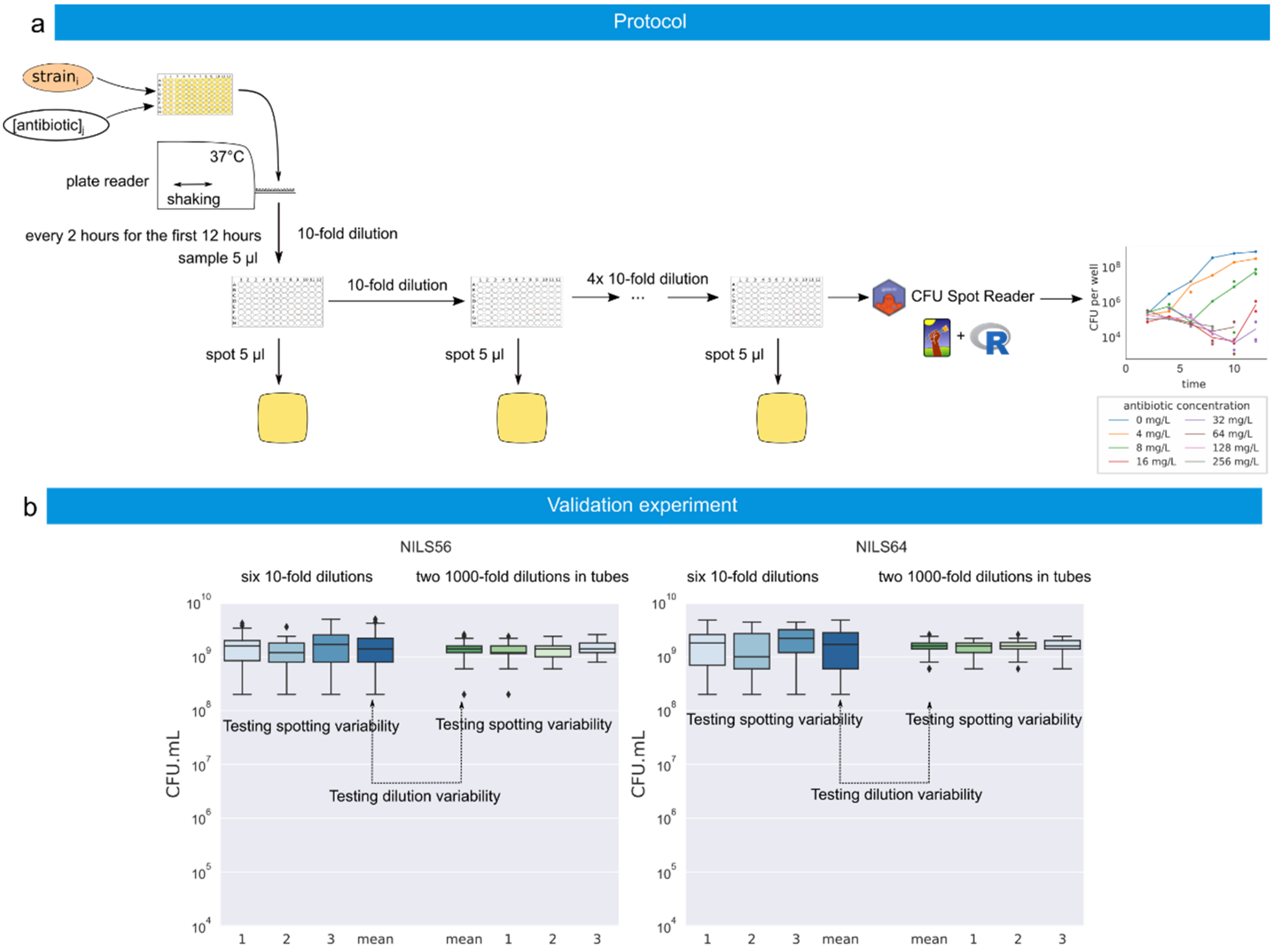

| strain | dilution type | CFU, mean | CFU, std | number of 10-fold dilutions | ratio | precision per dilution |
|---|---|---|---|---|---|---|
| NILS56 | six 10-fold dilutions | 1.68e+09 | 1.18e+09 | 6 | 1.19 | 0.97 |
| NILS56 | two 1000-fold dilutions in tubes | 1.41e+09 | 4.69e+08 | 0 | | |
| NILS64 | six 10-fold dilutions | 1.88e+09 | 1.35e+09 | 6 | 1.17 | 0.97 |
| NILS64 | two 1000-fold dilutions in tubes | 1.60e+09 | 4.40e+08 | 0 | | |

**Supplementary Figure 4.1 Presentation and validation of droplet-based CFU assay protocol.** a. *Graphical representation of the protocol.* Experimental plate containing bacterial culture, subjected to a range of antibiotic treatments, is put into a Tecan Spark plate-reader, where periodic OD measurements are taken and the plate is shaken and incubated at 37˚C. Every two hours plate is taken out of plate-reader to sample 5 µL and then returned to incubation and OD measurements. This sample is diluted tenfold six times. 5 µL of each dilution is spotted on agar-filled square Petri dish. Agar-filled plates are photographed after incubation. Images are analyzed using the CFU Spot Reader application. b. *Testing variabilities and validation experiment.* For two strains (NILS56 and NILS64) two different dilution protocols were performed: one just as described before (in blue), the other one consisted in doing two thousandfold dilutions in tubes (in green). In order to test spotting variability, last dilution in all four cases was spotted three times on agar plates (light shades). In order to evaluate dilution variability, we compared average values of all the results obtained using each dilution protocol (dark blue vs dark green). In the table we presented the mean value of CFU/mL for each case, ratio between the two means corresponding to different dilution protocols and $\sqrt[6]{\text{ratio}}$ to assess for precision per dilution.

**Supplementary Text 5: Model calibration on OD and CFU measures for antibiotic response characterization of different clinical isolates**

The whole strain collection presented in Supplementary Text 1 was submitted to a range of concentrations of cefotaxime for an in-depth characterization. For each strain and each antibiotic concentration, we measured the temporal evolution of both OD and CFU, and applied the two-step model calibration process described in the main text of this study. Here we present the results for all eleven strains, ranging from very susceptible to highly resistant to cefotaxime. Except for a few exceptions, discussed in the main text, model simulations are in good agreement with experimental data.

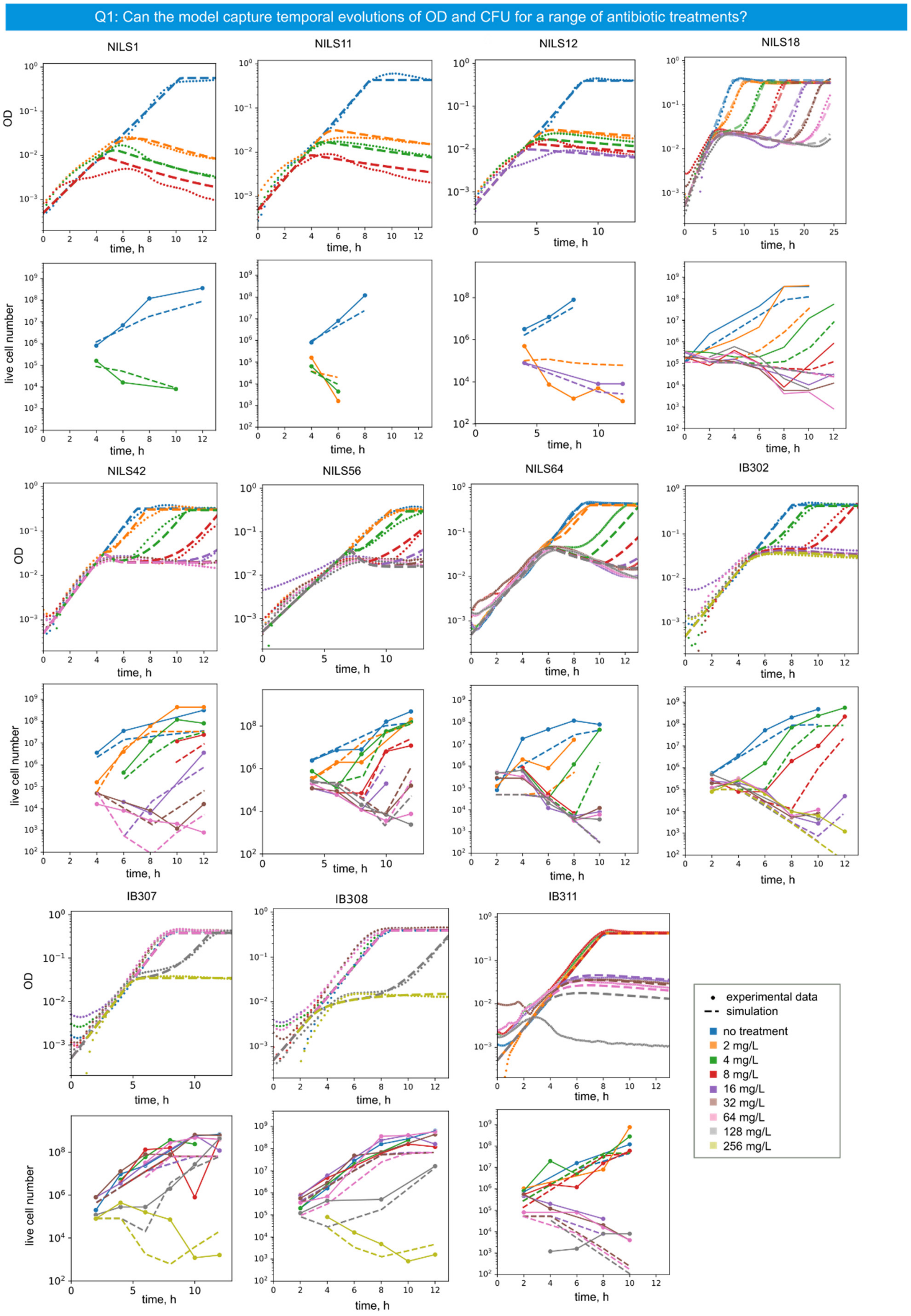

**Supplementary Figure 5.1 Comparison of experimental data and model simulations**. Eleven clinical isolates, representing a range of susceptibility to β-lactam treatment from very susceptible (NILS1, 11, 12), to highly resistant (IB307, IB308), are treated with different concentrations of cefotaxime, and optical density and live cell number are measured as described in Methods section.

We show in points with (CFU) or without (OD) solid lines the experimental data and in dashed lines the output of the model fitted on OD and CFU data simultaneously.

**Supplementary Text 6: The impact of genetic background on the response to antibiotic**

The non-susceptible strains, presented earlier in Supplementary Text 6, express a variety of different β-lactamases and represent different levels of antibiotic escape. In the interest of quantifying differences in phenotypes, we analyzed some of the parameter values from model calibration on OD and CFU data for these 8 strains (Supplementary Figure 5.1). For each strain, we selected the 45 best ones based on their fitting scores. Firstly, we looked into $k_1$parameter, which is the minimal concentration at which cells stop growing and dividing normally and start filamenting. The higher this value, the more resistant the strain is at the single cell level. On the left boxplot, three strains stand out due to high $k_1$ value: IB307, IB308 and IB311. This high value means that these strains can grow and divide normally subjected to higher concentrations of antibiotic than the other strains. This is confirmed by the experimental data. The strain with the lowest $k_1$ value is NILS31, for which we observe growth arrest even at smaller concentration of antibiotic. Secondly, we looked into $k_b * B_{in}$, a production of two parameters $k_b$ and $B_{in}$ which correspond to the degradation activity rate of the β-lactamases and the number of enzymes produced by unit of cell respectively. These two parameters form a structurally unidentifiable pair. For this reason, here, we consider the product of the two parameters. Interestingly, the three strains with highest $k_1$value have the lowest $k_b * B_{in}$values, especially IB311. This means, that these cells can grow normally until some relatively high concentrations, however, when subjected to even higher concentrations, the β-lactamases they express are not sufficiently efficient. In other words, these strains are individually resistant, but not very resilient as population. For IB311, this translates into a specific phenotype: at concentrations up to 8 mg/L there is no impact of antibiotic treatment on the growth dynamics, at one specific concentration from the tested range (16 mg/L), there is the phenomenon of crash, growth arrest and regrowth, and at higher concentrations (starting from 32 mg/L), there is crash and growth arrest, but no regrowth. This also concurs with the fact that one common attribute of all strains but this one is that they express a β-lactamase of CTX-M class.

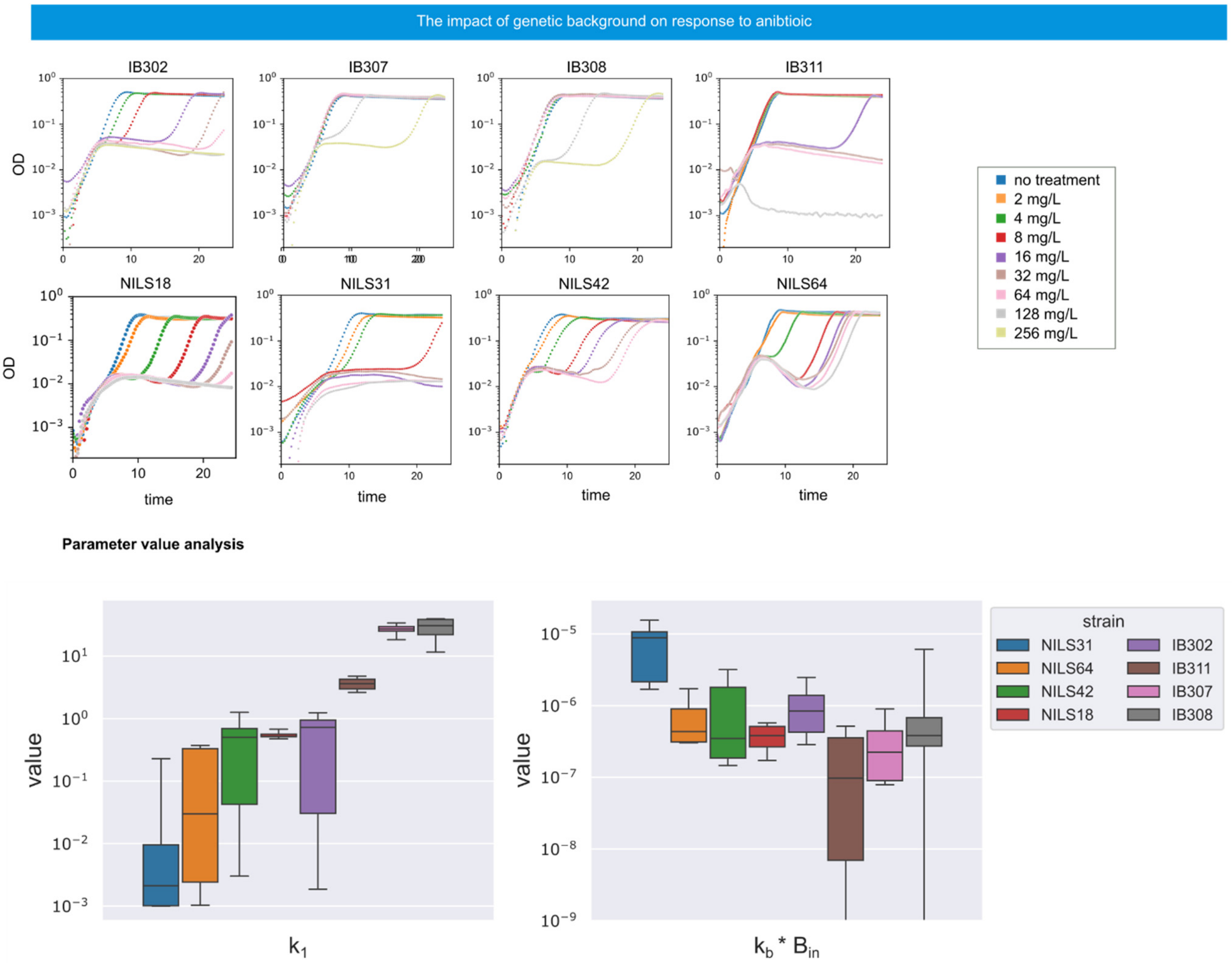

**Supplementary Figure 6.1 The impact of genetic background on bacterial population's response to antibiotic.** *Top:* Overview of bacterial population response to 8 cefotaxime concentration for 8 non-susceptible strains from the collection. *Bottom:* Boxplot of parameter values for $k_1$ (antibiotic concentration at which filamentation starts) and for $k_b * B_{in}$ (antibiotic degradation activity and production rate per cell length respectively) for 45 fits on OD and CFU for the 8 non-susceptible strains.

### Supplementary Text 7: Prediction of temporal evolution of CFU based on temporal evolution of OD

For five selected non-susceptible strains from the collection, we generated experimental data corresponding to single and repeated treatment experiments. For each strain we generated around 40 parameter value sets using only single treatment as calibration dataset and another 40 using single and three repeated treatment experiments as calibration dataset. For each parameter value set we calculated the CFU prediction score. The distribution of these scores is represented on histograms on the right of Figure 8.1. For most of the strains we clearly observe better scores while including repeated treatment experiments in calibration dataset. For others, these fits have good scores, but there is no clear separation between the two calibration datasets. On the same figure, we presented for each strain the four experiments composing the calibration dataset and the CFU simulations generated by the parameter value set having the median prediction score. Except for some high antibiotic concentrations, where there is some decorrelation between OD and CFU, these representative simulations are in good agreement with the experimental data.

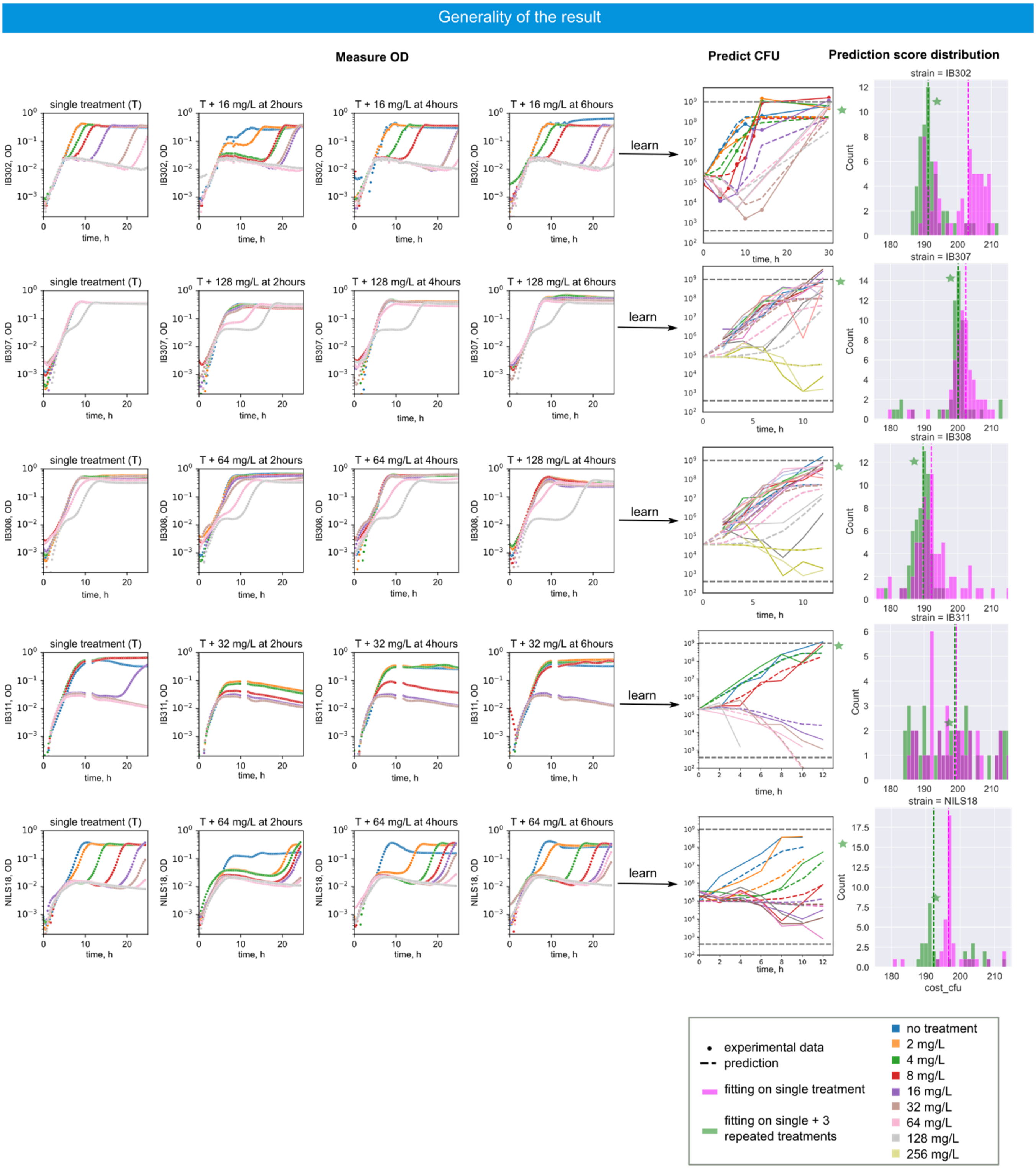

**Supplementary Figure 7.1 Prediction of temporal evolution of CFU based on temporal evolution of OD.** Each row of plots corresponds to one clinical isolate First plot in a row represents bacterial response to different antibiotic concentrations applied at the beginning (single treatment). Next three plots correspond to repeated treatments: to the range of initial single treatments a second antibiotic concentration is administered after several hours. Next plot shows experimental CFU data for single treatment (in points and solid lines) and the simulations (in dashed lines) using the parameter value set that corresponds to mean prediction score. The last plot in a row shows distribution of CFU prediction scores of different parameter value set that were calibrated either only on single treatment (in magenta) or on all four experiments (in green).

### Supplementary Text 8: Fitting on single treatments might lead to overfitting the data

In the main text, for one of the strains (IB302), we presented CFU predictions generated with parameter value sets calibrated on a single treatment experiment taken either alone or together with three repeated treatment experiments. We show representative predictions, we showed the predictions obtained by choosing the parameter set having the median CFU prediction score. Naturally, in a standard prediction problem, the CFU values are not available, so our proposed approach is not applicable. One might then wonder whether the parameter sets having the best prediction scores (on CFU) are those that have the best fitting scores (on OD). On Supplementary Figure 8.1, for each of the two calibration datasets, we selected the four best sets and generated the corresponding CFU predictions. We observed that although the fitting cost on the single treatment experiment alone is much lower than on the more complex calibration dataset (5 to 10 times), the CFU prediction score is lower when repeated treatment experiments are used for calibration. This leads us to think that given only single treatment experiment, parameter search tends to overfit the OD data and overestimate cell death and de-filamentation rate.

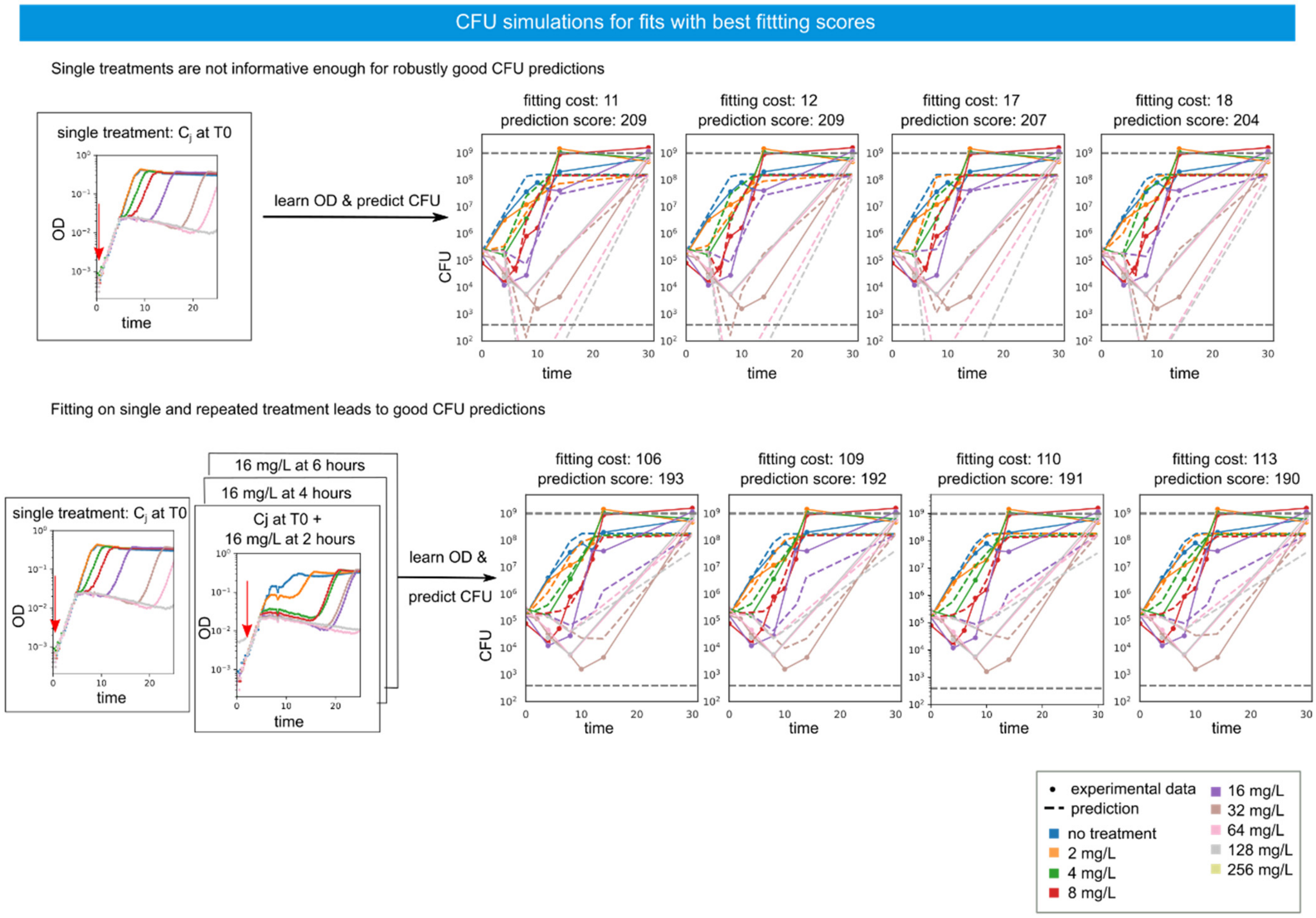

**Supplementary Figure 8.1 Prediction of temporal evolution of CFU using parameter values with lowest fitting score.** *Top:* Four parameter value sets, obtained through calibration on single treatment experiment and giving lowest fitting score on OD, were used to simulate temporal evolution of CFU. Each plot on the right corresponds to simulations from one parameter value set. In the title of each plot OD fitting score and CFU prediction score are presented. *Bottom:* Four parameter value sets, obtained through calibration on single and three repeated treatment experiments (represented on the left) and giving lowest fitting score on OD, were used to simulate temporal evolution of CFU. Each plot on the right corresponds to simulations from one parameter value set. In the title of each plot OD fitting score and CFU prediction score are presented.

### Supplementary Text 9: Impact of repeated treatment experiments on parameter search efficiency

Previously we compared the predictive power, defined in terms of CFU prediction score, of parameter value sets resulting from model calibration performed on a single treatment experiment taken either alone or together with three repeated treatment experiments. For Figure 3c and Supplementary Text 7, we present prediction results based on specific sets of three repeated treatment experiments. In most cases, a second antibiotic concentration has been applied at 2, 4 and 6 hours after the first dose. Therefore, one might wonder how many additional data sets are needed to obtain good CFU prediction scores. One might also wonder whether only their number matters or whether some repeated treatments are more informative than others. Lastly, to what degree are parameter values better constrained when we add datasets in the calibration data?

To investigate this question, we have performed 7 different repeated treatments for two strains (IB302 and IB311) and 3 different repeated treatments for three other strains (IB307, IB308, and NILS18). We then generated different calibration datasets, called dataset types, containing the single treatment experiment and a variable number of repeated treatments. For IB302 for example, taking the single treatment experiment and one of the seven repeated treatment experiments leads to a dataset of type “mono+1r2”, and adding to these datasets a second repeated treatment experiment leads to a dataset of type “mono+2r2”. There are 7 variants of the first type and 21 variants of the second type. For all possible dataset types, and all possible combination of experiments in each given type (ie, for all variants of the type), we repeatedly calibrated the model on the considered dataset. The number of dataset types, of dataset variants in a given type, and of successful calibration results is summarized in the table provided on Supplementary Figure 9.1 (top, left). For each fit, we calculated the fitting score on the OD of the calibration dataset and the prediction score on the CFU of the single treatment experiment. Based on previous observations and analysis, we have set 200 as a threshold for both scores to separate good fits from medium or even bad ones. We show for three strains (IB302, IB307 and NILS18) and for each dataset type the percentage of fits having a good fitting score and the percentage of fits having a good prediction score among the fits having good fitting score (Supplementary Figure 9.1; top, middle). From these plots, we can observe that adding repeated treatment experiments tends to decrease the probability to have a good fit, however, it increases chances of a fit with good fitting score to have a good prediction score. In other words, it is harder to fit on more complex dataset, but if we get a good fit, it is more likely that it will generate good CFU predictions. In order to consolidate this result, we looked into the evolution of fitting scores and prediction scores as a function of the number of experiments in the calibration dataset (Supplementary Figure 9.1; top, right). We observe that the average fitting score goes up, when we add at least one repeated treatment experiment, and, at the same time, if we compare fits on only single treatment experiment and fits on single and three repeated treatment experiments, we observe that, in the latter case, the average prediction score is lower, and, importantly, many variants in this dataset lead to very good prediction scores (below 190).

In order to study the impact of the choice of a specific calibration dataset on the uncertainty of the parameter values, we calculated for each dataset variant the standard deviation of the set of parameter values obtained after calibration. On Supplementary Figure 9.1 (bottom), we show this data for a small selection of parameters: growth rate $\mu$, a parameter easily identifiable with minimal amount of data; antibiotic concentration inducing filamentation $k_1$, an antibiotic-related parameter, important for good CFU predictions, but still identifiable from single treatment experiment; and, finally, the product of two parameters characterizing production and efficiency of β-lactamase $k_b * B_{in}$, hard to identify, but important for good CFU predictions. As expected, the growth rate is well constrained by the data on all calibration datasets. For IB302, but also IB307, we observe that adding experiments tends to add constraints to parameter values. For NILS18, this effect depends a lot on which three repeated treatment experiments is chosen.

Overall, choosing as calibration dataset a single treatment experiment together with three repeated treatment experiments  is a good compromise between the complexity of the calibration dataset and the quality of the CFU predictions and the identifiability of model parameters.

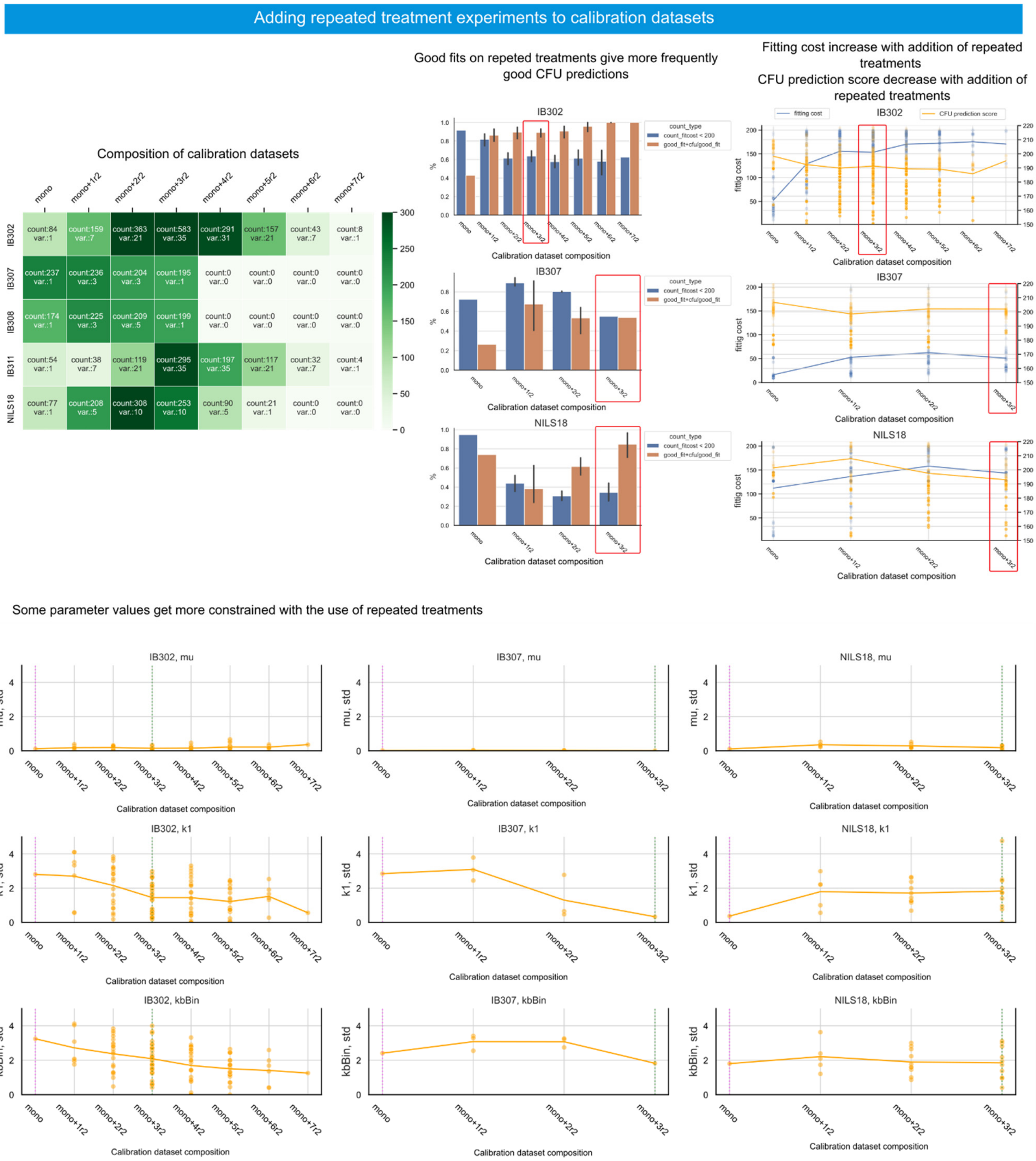

**Supplementary Figure 9.1 Impact of addition of repeated treatments to the calibration dataset on parameter fitting**. *Top:* Analysis of fit quality depending on calibration dataset type. *Left:* This table represents a summary of number of generated fits for each strain and for each dataset type. “Count” and color of the cell correspond to total number, “var.” corresponds to number of individual variants of this type with at least one fit. *Center:* For each individual dataset total number of fits, number of fits with fitting score less than 200 and number of fits with both prediction score and fitting score are less or equal to 200 were calculated. On this plot blue bars correspond to average per dataset type percentage of fits with good fitting score, orange bars correspond to average per dataset type percentage of fits with good prediction score among those with good fitting score. Error bars represent variability between different individual datasets of the same type. *Right:* This plot shows evolution of fitting and prediction scores as a function of calibration dataset type. Points represent individual parameter value sets; solid lines represent average per dataset type. Blue color corresponds to fitting score, orange color corresponds to CFU prediction score. *Bottom:* Analysis of parameter value constraints. For each individual dataset for each parameter a standard deviation of value distribution was calculated, represented here in dots. Solid lines correspond to average per dataset type. Different columns correspond to different clinical isolates. Different rows correspond to different parameters. Here we selected three parameters: growth rate $\mu$ (first row), a parameter easily identifiable with minimal amount of data; antibiotic concentration inducing filamentation $k_1$ (second row), a antibiotic-related parameter, important for good CFU predictions, but still identifiable from single treatment experiment; and, finally, product of two parameter characterizing production and efficiency of β-lactamase $k_b * B_{in}$ (third row), hard to identify, but important for good CFU predictions.

#### Supplementary Text 10: Predicting complex temporal evolution of CFU for delayed treatments

As an additional challenge for the predictive power of the model, we sought to predict the temporal evolution of CFU for delayed treatments: the entire antibiotic dose is administered several hours after the beginning of the experiment. This is a version of the inoculum effect since the bacterial population has time to grow to medium or high densities before antibiotic administration. The connection between OD and CFU data is even more complex for delayed treatments than it was for initial treatments. For instance, there is no difference in temporal evolution of OD between treating at 4 hours or at 6 hours or normal growth with no treatment. However, the evolution of the CFUs is drastically different in these cases.

On Figure 4a we showed experimental data and model simulations for several delayed treatments with one antibiotic concentration (16 mg/L). Here we present similar results for another antibiotic concentration (32 mg/L). Just as before, we used the same parameter value set as on Figure 3, that was obtained through calibration on OD data of one single and three repeated treatments experiments. Note that our calibration dataset contained the OD information corresponding to (0 mg/L of cefotaxime at time 0 and) 16 mg/L of cefotaxime added at either 2, 4 or 6 hours. The OD, but not the CFU data, for delayed treatments with 16 mg/L was present in the calibration data set. However, neither the OD or the CFU data for delayed treatments with 32 mg/L of cefotaxime were present in the calibration dataset. This could  explain a slightly larger difference between experiments and simulations in the latter case. Overall, the simulations are in good agreement with the data, especially considering the complexity of the data.

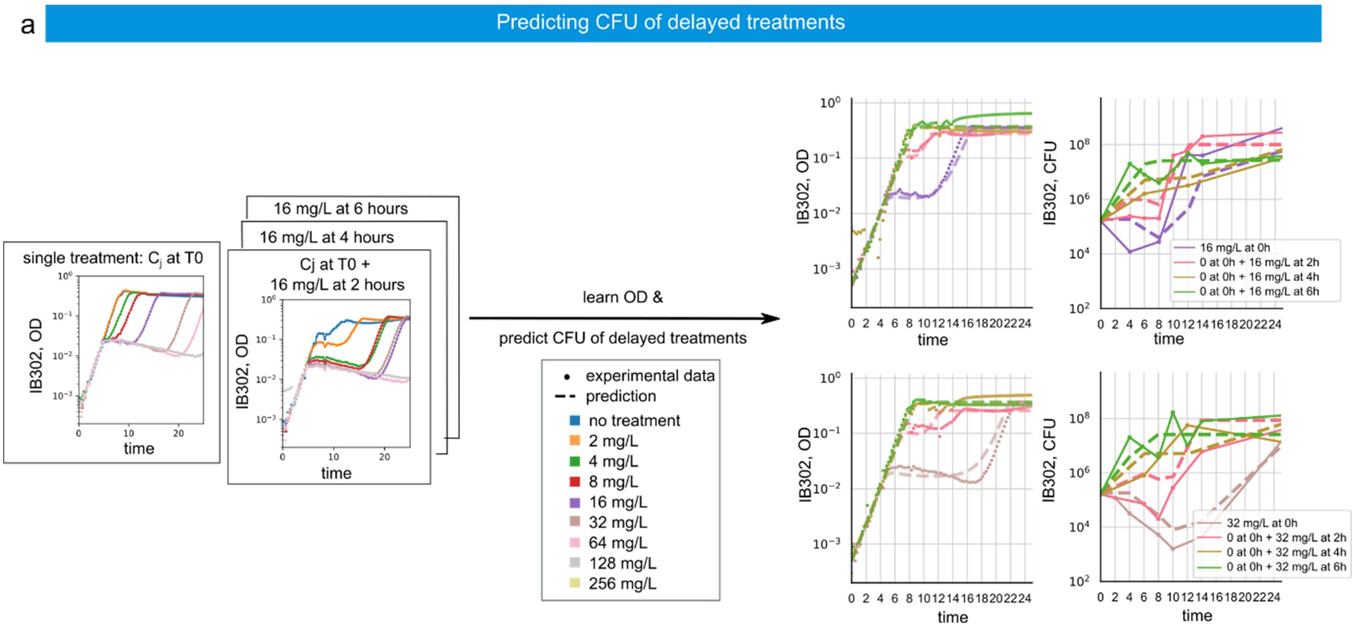

**Supplementary Figure 1.1 Predicting the complex temporal evolution of CFU data for delayed treatments**. The calibration dataset is represented on the left side. Experiments corresponding to the data on the plots on the right side are delayed treatment experiments, where there is no antibiotic added at the beginning of experiment, and a given concentration is administered several hours later. The top row corresponds to treatments with 16 mg/L of cefotaxime; the bottom row corresponds to 32 mg/L. Experimental data is presented in points with solid lines and simulations are represented by dashed lines.

#### Supplementary Text 11: Optimal treatments leveraging glucose exhaustion

In the main text we presented an optimal treatment solution for IB311. Here, we consider NILS18. As before, we fitted the model using as calibration dataset one single and three repeated treatments experiments. The optimal treatment that minimizes the final OD at thirty hours was predicted to be the addition of the entire dose 3 hours after the beginning of the experiment. This might be explained by the consumption of nutrients by cells that will die upon antibiotic treatment, leading to glucose depletion and incomplete full regrowth. This scenario of final OD of delayed treatment being lower than the carrying capacity is confirmed by experimental data. On the Supplementary Figure 11.1, we present experimental data and corresponding predicted behaviors for the predicted optimal treatment, and close alternative treatments.

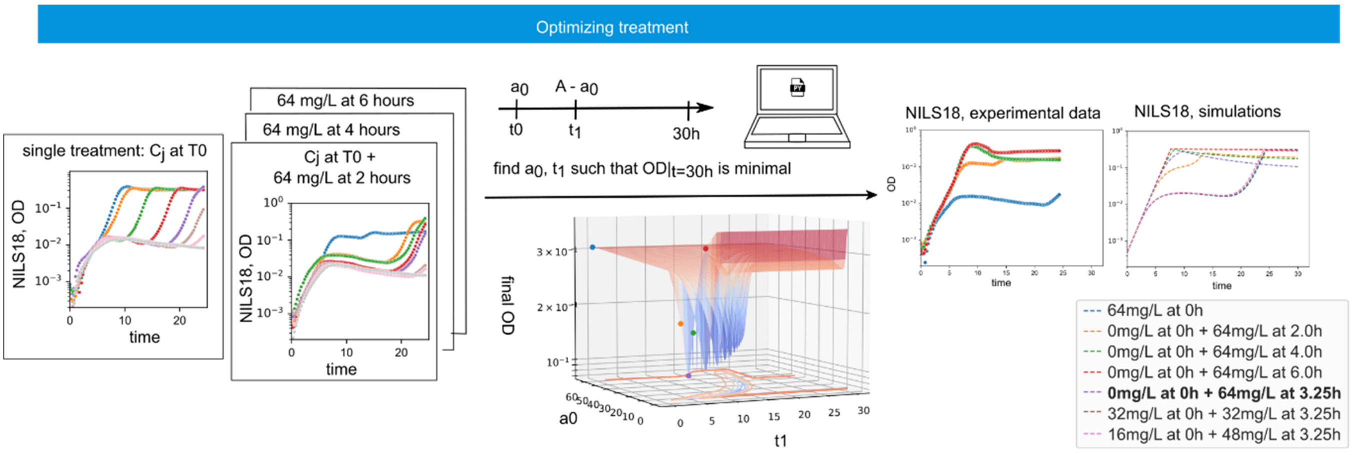

**Supplementary Figure 11.1 Optimal treatment through glucose exhaustion**. *Optimization problem:* we search for the most efficient way to split a given antibiotic concentration into two separate doses, one ($a_0$) at the beginning and the rest at time $t_1$ so that OD at 30 hours is minimal. *Left*: calibration dataset and fitting results. *Center:* surface of the optimization search in the optimization space. *Right:* Experimental and *in silico* results. The position of the tested treatments are shown on the optimization surface by the points of the corresponding colors.
